# Fibroblast-derived Collagen VI shapes the structure and function of the tumor-immune microenvironment in clear cell renal cell carcinoma

**DOI:** 10.64898/2026.03.19.712351

**Authors:** Maximilian Wess, Grigor Andreev, Tobias Feilen, Nastasja Diel, Max Zirngibl, Constance Gueib-Picard, Anna L. Kössinger, Célia Hinrichs, Andreas Nägel, Clara L. Essmann, Winfried Römer, Martin Helmstädter, Tino Vollmer, Martin Werner, Markus Grabbert, Oliver Schilling, Manuel Rogg, Christoph Schell

**Author notes:** contributed equally (Manuel Rogg, Christoph Schell). To whom correspondence should be addressed: Prof. Dr. Dr. Christoph Schell Institute of Surgical Pathology University Medical Center Freiburg; Breisacherstr. 115a, D-79106 Freiburg, Germany.

## Abstract

The importance of the extracellular matrix (ECM) influencing tumor biology in stroma-rich tumors is well established. However, the relevance of individual ECM proteins in rather stroma-poor cancers such as clear cell renal cell carcinoma (ccRCC) is ill-defined. Using bulk proteomics, spatial imaging, and single-cell transcriptomics, we identify collagen VI (COL6) as a predominant ECM component of the ccRCC interstitial stroma, synthesized primarily by fibroblasts and pericytes. Using cell-derived matrix (CDM) models, we demonstrate that COL6 is essential for maintaining an isotropic ECM network architecture and governs the broader matrisomal composition, with direct pro-proliferative consequences for tumor cells both in vitro and in situ. Granular spatial analysis reveals that COL6-rich stromal septa constrain tumor-infiltrating T cells to boundary zones, where CD8+PD1+ phenotypes predominate. Importantly, tyrosine kinase inhibition (TKI) with cabozantinib suppresses COL6 expression in fibroblasts in vitro and in ex vivo tumor models, mirroring COL6-depleted CDM phenotypes. Our findings establish COL6 as a central stromal regulator of ccRCC tumor biology and immune contexture, revealing ECM remodeling as an underappreciated mechanism of TKI action, with implications for combination immunotherapy strategies.

## Introduction

Bidirectional cell-matrix interactions play an essential role in a plethora of physiological processes and are critically involved in cancer progression^1, 2^. The composition and mechanical properties of tumor-associated extracellular matrix (ECM) significantly influence the functional architecture of the tumor-immune-microenvironment (TIME) and thus the biology of respective tumors. In this context, stroma/ECM-rich tumors often exhibit a more aggressive biological behavior characterized by increased invasive growth, metastasis and therapy resistance^2, 3^. Interestingly, even in rather stroma-poor tumors such as clear cell renal cell carcinoma (ccRCC), the most prevalent type of renal cancer, predictive ECM-signatures have been identified^4, 5^. However, a detailed mechanistic understanding of how individual ECM proteins potentially orchestrate the functionality of the TIME in ccRCC is missing.

Aside from its name-giving “clear cell appearance” (due to the high content of intracellular lipids), ccRCC shows an expansive growth pattern resulting in the formation of a pseudocapsule at the tumor-boundary zone^6, 7^. Given the highly prevalent mutational loss-of-function of *VHL* and consequently increased pseudohypoxic signaling (due to accumulation of HIF1/2 transcription factors), ccRCC is also characterized by a high degree of vascularization^8, 9^. Therefore, the histological architecture of ccRCC is mainly determined by a dense accumulation of tumor cells and rather minute amounts of stromal septa, where fibroblasts, pericytes, endothelial cells and immune cells are primarily detected. Although the stromal/ECM content is obviously minor when compared to other solid tumors such as pancreatic adenocarcinoma, colorectal and ovarian cancer^10^, recent studies have identified specific ECM-signatures corresponding to aggressive biological behavior^6, 11^, immune modulation^7, 12^ and consecutive therapeutic implications^13^. Interestingly, the majority of studies focused so far on the role of respective ECM-proteins directly within the tumor cell population^14^, and did not address the predominant ECM-producing niche, which is constituted by the perivascular stroma.

In ccRCC, the majority of deposited ECM is detected in very thin fibrovascular septa embedded between clusters of tumor cells. This compartmentalized histoarchitecture results in close spatial proximity between deposited ECM and remaining cellular populations within the TIME niche. Following a proteomic-based approach to annotate the general ECM repertoire within ccRCC, combined with further spatial imaging approaches, led to the identification of collagen VI (COL6) as one predominantly expressed ECM protein in the interstitial compartment of ccRCC. COL6 is a ubiquitously expressed member of the large collagen-protein family, contributes to filamentous ECM networks (fibrillar collagen) and exhibits a heterotrimeric microfibril structure mainly composed of three COL6-α chains (COL6A1, COL6A2 and COL6A3)^15^. Based on the large variety of structural domains, COL6 interacts and binds to numerous components of the ECM, thereby promoting cell-matrix interactions and organizing various tissue architectures^16^. In cancer, COL6 expression appears to be tumor-type dependent^17^, but it frequently propagates tumor growth, progression, and metastasis^18^. Aside from direct interaction with chondroitin sulfate proteoglycan receptors and consecutive AKT/β-catenin signaling^19, 20^, a C-terminal cleavage product (endotrophin – ETP) has been implicated in TGFβ signaling as well as induction of epithelial-mesenchymal transition (EMT)^21, 22, 23^. In the context of ccRCC, recent studies have focused on the direct impact of COL6 within respective tumor cells. For example, it has been observed that COL6A2 was linked to stem cell-like properties of cancer cells and might regulate processes such as epithelial-mesenchymal transition via PI3K-Akt signaling^24^.

Here, we demonstrate that COL6 is predominantly localized within the interstitial stroma of ccRCC and that COL6 influences the cell-ECM interface with differential impact on cellular dynamics of fibroblast, immune and tumor cells. Importantly, we show that COL6-depletion not only profoundly alters the structural architecture of the matrix but also results in major compositional changes of the ECM. We further demonstrate that these changes translate into functional consequences for immune and tumor cell populations, which are also transferable to the *in situ* context. We further observe that commonly employed tyrosine kinase inhibition (TKI) directly regulates COL6 deposition within the tumor stroma. Therefore, we propose COL6 as a critical regulator of tumor-stroma dynamics with potential therapeutic implications in ccRCC.

## Results

### COL6 is overexpressed in ccRCC and predominantly localized in the tumor stroma

To annotate the global ECM composition in ccRCC, we employed a multilayered analysis approach involving bulk tissue-proteomics, histology-based machine learning and multiplex immunofluorescence (Figure 1A). Proteomic analysis on a ccRCC patient cohort (N=23 tumors) indicated a pronounced abundance of COL6 within the tumor tissue (including its three predominant chains, COL6A1, -A2 and -A3) when compared to matching normal adjacent tissue (NAT, N=18), in line with an overall accumulation of matrisomal proteins (Figure 1B&C). GO enrichment analysis further demonstrated a distinct matrisomal pattern in tumor tissue of ccRCC characterized by enrichment of collagen fibril organization and collagen-containing ECM (Figure 1D). Interestingly, individual COL6 chains appear to be almost exclusively upregulated in ccRCC, indicating a potentially decisive role for COL6 in the tumor immune microenvironment of ccRCC (Figure 1C). This is further supported by observations of impaired overall survival of patients with high expression levels of COL6A1, COL6A2 and COL6A3 (Supplementary Figure S1). Employing a machine learning based segmentation approach for classification and quantification of tissue compartments indicated a positive correlation between COL6 protein expression and the extent of the tumor stroma (Figure 1E&F). Further granular analysis showed that these correlations do not apply to segmented mesenchymal cells, but rather the stromal compartment in line with deposited ECM (Figure 1F-H and Supplementary Figure S1). Leveraging single-cell transcriptomic analysis of established ccRCC cohorts correspondingly demonstrates the predominant synthesis of COL6 by fibroblasts and pericytes (Figure 1I and Supplementary Figure S1). Together, these findings collectively highlight that COL6 is predominantly deposited in the tumor stroma and synthesized by fibroblasts, where it constitutes a major ECM component potentially influencing the tumor microenvironment and biology of ccRCC.

**Figure 1.**
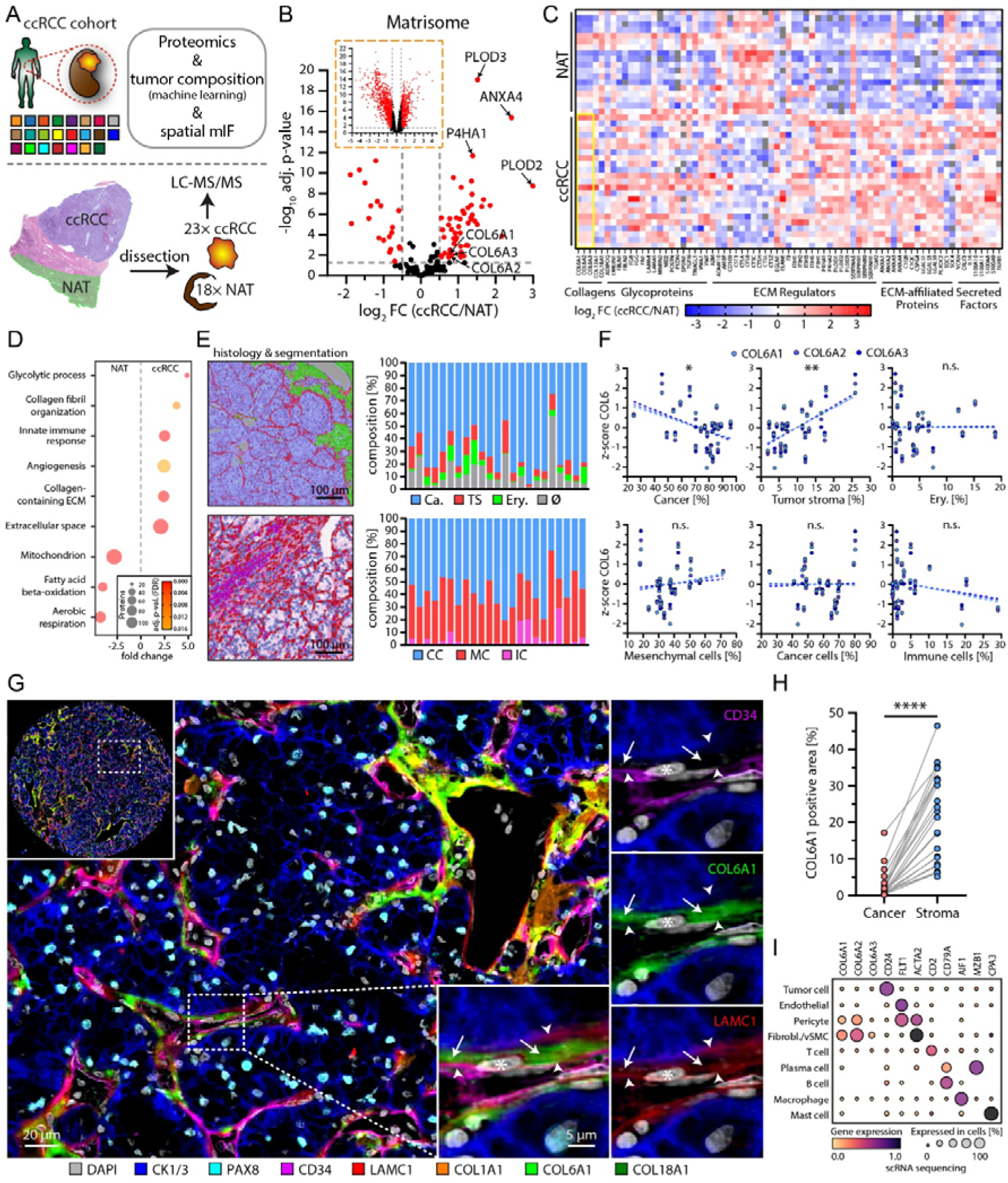
COL6 is overexpressed in ccRCC and predominantly localized in the tumor stroma. **(A)** Schematic depicting the processing of a cohort of cases of 23 ccRCCs for proteomic analysis, histological tumor segmentation and spatial multiplex immunofluorescence (NAT – normal adjacent tissue). **(B)** Volcano plot analysis of the ccRCC proteome highlighting significantly regulated matrisome proteins (red dots indicate proteins with an adjusted p-value <0.05 and log_2_ FC (fold change) > |0.5|; dashed boxed show unfiltered proteomic analysis). **(C)** Heatmap presentation of significant differentially expressed matrisome proteins in the proteomic analysis of ccRCC samples sorted by matrisome categories (yellow box highlights upregulated collagen-6 chains in ccRCC tissue). **(D)** GO enrichment analysis shows an increase in categories such as collagen fibril organization in ccRCC tumor tissue. **(E)** Compartment segmentation of ccRCC tumor tissue (top – Ca. = cancer (blue), TS = tumor stroma (red), Ery. = erythrocytes (green), Ø = no tissue (cysts, vessels) (grey)) and cells (bottom – CC = cancer cells (blue), MC = mesenchymal cells (red), IC = immune cells (violet)). **(F)** Sample wise correlation analysis of histological compartment composition by machine learning based segmentation shown in panel (E) and z-scores of COL6A1, COL6A2 and COL6A3 expression in the ccRCC proteome (N=23 patients, Pearson correlation; _∗_ – p < 0.05, _∗∗_ – p < 0.01 and non-significant (n. s.)). **(G)** Multiplex IF staining demonstrates predominant deposition of COL6 in the interstitial compartment (dashed boxes indicate regions of magnified areas, asterisk indicates perivascular mesenchymal cells (Fibroblasts, Pericytes), arrows indicate the ECM, arrowheads indicate the basement membrane). **(H)** Quantification of COL6A1 positive area in cancer and stroma tissue compartments of ccRCC (N=23 patients, _∗∗∗∗_ – p<0.0001, paired t test). **(I)** Dot plot heatmap of gene expression in indicated cell populations derived from available scRNA sequencing analysis of ccRCCs (N=7 patients).

### COL6 modulates the cell-matrix interaction by regulating focal adhesion formation and ECM gene expression

Given the predominant expression and localization of COL6 within the tumor stroma of ccRCC (Figure 1, Supplementary Figure S1 and S2), we aimed to differentiate between potential effects of COL6 on tumor cells via extrinsic interaction (via the ECM) and potential intrinsic effects^24^. First, we generated a knockdown of *COL6* employing shRNA approaches on the background of the 786-O ccRCC cancer cell line (Figure 2A and Supplementary Figure S2). Here, no direct impact on tumor cell proliferation was evident (based on colony formation and BrdU incorporation assays; Figure 2 B&C). Interestingly, we observed formation of larger colonies in respective knockdown cells, while the colony number (and therefore clonogenic potential) was not affected (Figure 2B). This altered colony morphology might be partially explained by direct effects of *COL6* knockdown on cellular features such as focal adhesion formation (Figure 2D) and transcriptional alterations involving pathways such as cell-substrate interaction (Figure 2E). Interestingly, acute consequences of this impeded cell-matrix interaction do not directly affect cellular functions such as collective cell migration (Supplementary Figure S2). Based on the distinct localization pattern of COL6 within the stromal compartment of the tumor septa in ccRCC (Figure 1), we primarily focused on the role of COL6 in fibroblast populations (Figure 2F). While not reaching statistical significance, depletion of COL6 in the renal fibroblast cell line TK173 translated into a similar colony morphology as observed in cancer cells but did not directly affect proliferative behavior (Figure 2G&H). Of note, a reverse experimental approach (overexpression of *COL6A2*) resulted in the opposite phenotype (Supplementary Figure S2). Moreover, in the fibroblast background, focal adhesion morphology and related transcriptional signatures were comparable to the observed effects in ccRCC cancer cells (Figure 2I&J). Notably, global comparison of transcriptional alterations between these two cellular backgrounds (786-O vs. TK173) indicates a more pronounced transcriptional response due to COL6 knockdown in fibroblasts compared to cancer cells (Figure 2E, 2J and Supplementary Figure S2B). This differential sensitivity likely reflects the higher basal COL6 expression levels in fibroblasts, which serve as the primary COL6-producing cells in the tumor microenvironment (Supplementary Spreadsheet S2). The greater transcriptional perturbation in fibroblasts upon COL6 depletion suggests that these cells experience more substantial alterations in cell-ECM adhesive signaling when COL6 is lost. Although COL6 itself is not a transcription factor, its depletion profoundly impacts integrin-mediated mechano-transduction pathways, which are potent regulators of gene expression. Together, our findings suggest that COL6 modulates the cell-ECM interaction independently of cell type (fibroblasts and cancer cells). Given the predominant extracellular deposition of COL6 and its established function as an integrin ligand, this supports a model in which COL6 acts as an external matrix cue that shapes focal adhesion morphology and ECM constitution to instruct cell-ECM interaction.

**Figure 2.**
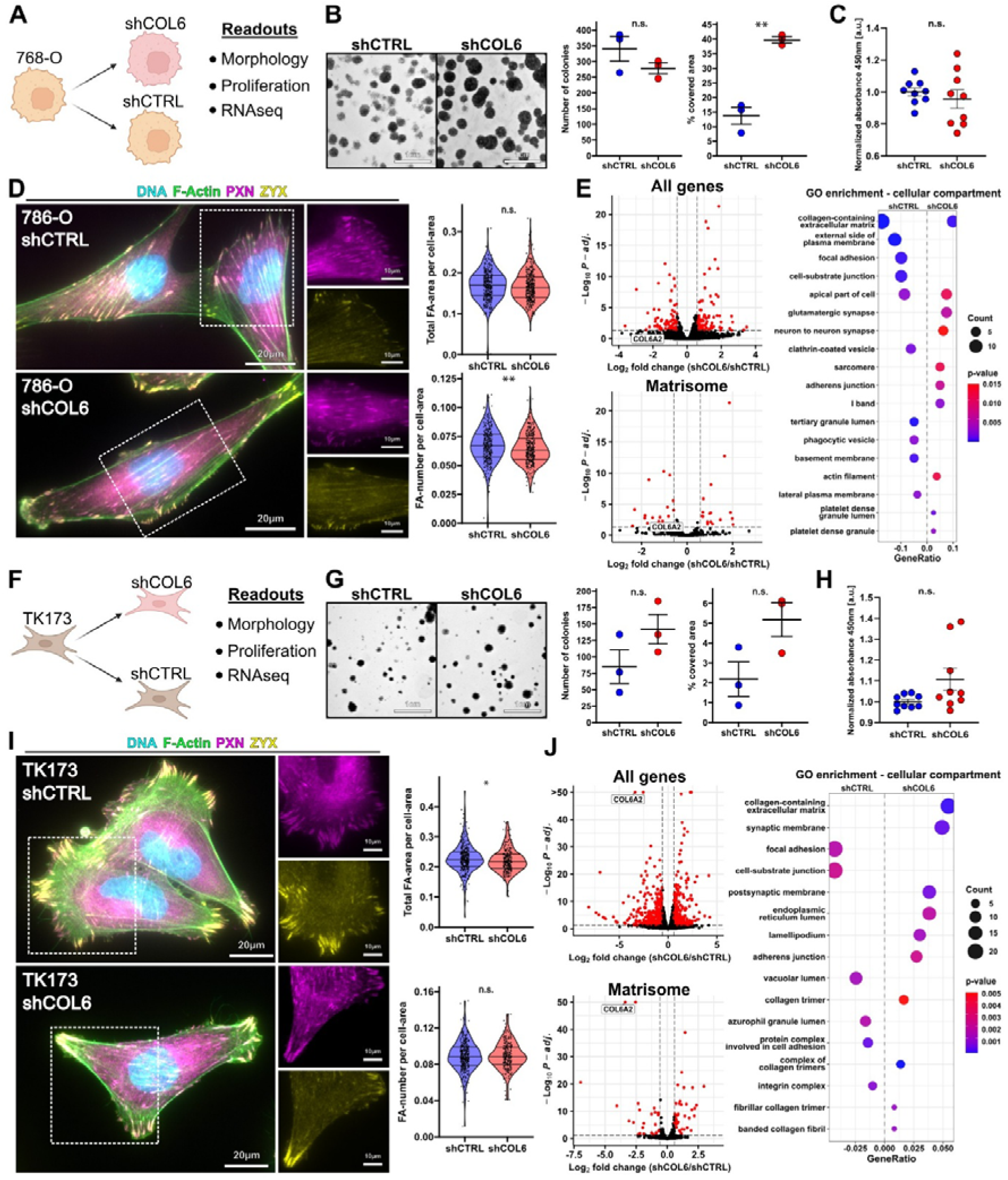
COL6 modulates the cell-matrix interaction by regulating focal adhesion formation and ECM gene expression. **(A)** Schematic representation of the generation of COL6 knockdown in the 786-O cell line. Created in BioRender. Schell, C. (2026) https://BioRender.com/x5sqb3y. **(B)** Representative images of colony formation assays of 786-O shCTRL and shCOL6 cells; Dot plots depict the number of colonies and the relative area covered (N=3 independent experiments, unpaired t test). **(C)** Dot plot indicating cell proliferation analysis by BrdU in 786-O shCOL6 and shCTRL cells (N=9 replicates of 3 independent experiments, performed in triplicate and pooled for analysis, unpaired t test). **(D)** IF staining of focal adhesion (FA) markers paxillin (PXN, purple) and zyxin (ZYX, yellow), as well as Nucleus (blue) and F-actin (green) in 786-O shCTRL and shCOL6 cells (white dashed boxes highlight area of zoom-in; Violin plots depicting FA area (top) and FA number normalized to respective cell area (dots indicate cells from 3 independent experiments, Mann-WhitneyLULtest). **(E)** Volcano plot of differential gene expression of all genes (top left) and matrisome-filtered genes (bottom left) in 786-O *COL6A2* knockdown and control cells; Right panel shows gene-ontology (GO) enrichment analysis represented in a bubble plot. **(F)** Schematic representation of the generation of COL6 knockdown in the TK173 cell line. Created in BioRender. Schell, C. (2026) https://BioRender.com/x5sqb3y. **(G)** Representative images of colony formation assays of TK173 shCTRL and shCOL6 cells; Dot plots depict the number of colonies and the relative area covered (N=3 independent experiments, unpaired t test). **(H)** Dot plot indicating cell proliferation analysis by BrdU in TK173 shCOL6 and shCTRL cells (N=9 replicates of 3 independent experiments, performed in triplicate and pooled for analysis, unpaired t test). **(I)** IF staining of focal adhesion (FA) markers paxillin (PXN, violet) and zyxin (ZYX, yellow), as well as Nucleus (blue) and F-actin (green) in TK173 shCTRL and shCOL6 cells (white dashed boxes highlight areas of zoom-in; Violin plots depicting FA area (top) and FA number normalized to respective cell area (dots indicate cells from 3 independent experiments, Mann-WhitneyLULtest). **(J)** Volcano plot of differential gene expression of all genes (top left) and matrisome-filtered genes (bottom left) in TK173 *COL6A2* knockdown and control cells; Right panel shows GO enrichment analysis represented in a bubble plot. Bars indicate mean and S.E.M in dot plots or median and quartiles in violin plots; ∗ – p < 0.05, ∗∗ – p < 0.01 and n.s – not significant.

### Depletion of COL6 translates into an anisotropic ECM architecture

To investigate the effects of COL6 on the biophysical and molecular properties of synthesized ECM, cell-derived matrices (CDMs) were employed as a model system^25^. CDMs were synthesized by control and *COL6* knockdown cells over a period of 7 days, followed by gentle decellularization and immunostaining, confirming efficient depletion of COL6 in knockdown conditions (Figure 3A-B). Interestingly, the loss of COL6 protein was accompanied by a significant alteration of matrix structure: fibers of the COL6-deficient CDM showed a more parallelized microarchitecture with less interfibrillar space compared to the stronger interconnected fibers in control conditions (Figure 3B and Supplementary Figure S3). This anisotropic matrix architecture in shCOL6-CDMs prominently evolves over time of CDM synthesis and matrix deposition (Supplementary Figure S3). The more parallelized formation of Fibronectin (FN) fibers in the COL6-deficient matrix is further reflected by an increase in equally distributed fiber orientations (Figure 3C). These observations imply that depletion of COL6 directly impacts the organization of the ECM architecture and also translates into a diminished thickness of the respective CDM (Figure 3D). Notably, knockdown of COL6 in 786-O cancer cells did not result in any detectable changes in the respective ECM structure (Supplementary Figure S3). This might relate to generally less matrix being deposited by 786-O and ccRCC cancer cells, or the initially rather low expression levels of COL6 in this cellular background (Supplementary Figure S2). On the contrary, neither the overexpression of COL6A2 in fibroblast populations resulted in an altered amount of COL6 being deposited nor did it translate into any structural alterations (Supplementary Figure S3). By using super-resolution microscopy (Airyscan), we further observed that COL6 localizes along FN fibers as well as in the interfibrillar space (Figure 3E), suggesting a modulatory effect of COL6 on fiber connectivity. To further visualize the ultrastructure of COL6-dependent ECM, we employed scanning electron microscopy (SEM). Here, we observed that upon depletion of COL6, the overall CDM structure appeared more fibrillar or fragmented in contrast to a more cohesive structure in COL6-containing CDMs (Figure 3F). As these structural changes within CDMs also potentially impact their biomechanical properties, we applied atomic force microscopy (AFM). However, AFM measurements showed comparable stiffness values (Youngśs Moduli in Pa) between both conditions (Figure 3G). To dissect whether structural alterations result from direct ECM effects or are indirect consequences of COL6 depletion (such as transcriptional regulation in fibroblasts; Figure 2J), we performed targeted rescue experiments. Remarkably, COL6 supplementation reversed the anisotropic ECM architecture and restored interconnected networks (Figure 3H, 3I and Supplementary Figure S3). The specificity of this effect was furthermore supported by supplementation experiments using COL1, where anisotropy of the deposited ECM was largely unaffected. Collectively, these observations establish COL6 as a central factor governing both matrix organization and surface ultrastructure. Having revealed COL6’s decisive role as a central matricellular organizer, we next sought to determine how its depletion reshapes the global compositional landscape of the matrisome (i.e., describing all components of ECM-proteins and related modifiers).

**Figure 3.**
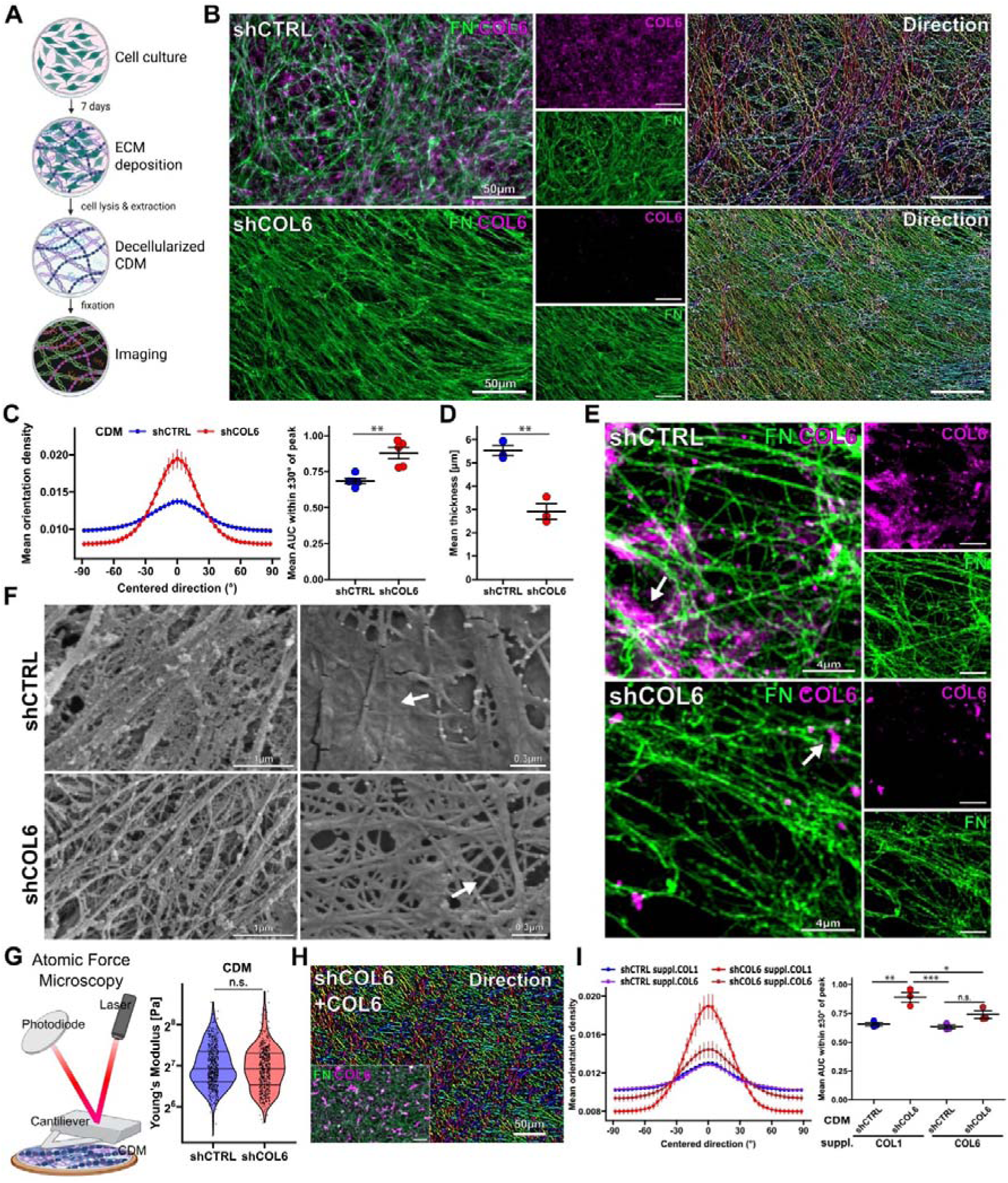
Depletion of COL6 translates into an anisotropic ECM architecture. **(A)** Schematic depicting the generation and imaging of cell-derived matrices (CDM). Created in BioRender. Schell, C. (2026) https://BioRender.com/x5sqb3y. **(B)** IF of CDM samples generated from TK173 shCTRL and shCOL6 cells stained for Fibronectin (FN, green), Collagen VI (COL6, violet) and color-coded FN-fiber direction. **(C)** Density plot (left) showing quantification (mean and S.E.M.) of FN-fiber orientation in shCTRL (blue) and shCOL6 (red). Dot plot (right) of mean area under curve (AUC) within ±30° centralized peak (N=5 independent experiments, unpaired t-test). **(D)** Dot plot of mean CDM thickness of CDMs derived from shCTRL and shCOL6 cell lines (N=3 independent experiments, unpaired t-test). **(E)** High-resolution IF imaging of CDMs from shCTRL (top) and shCOL6 (bottom) for FN (green) and COL6 (violet). White arrows indicate patch-like COL6 structures. **(F)** SEM imaging of CDMs with an overview (left) of shCTRL and shCOL6 and zoom-ins (right). The white arrow in the shCTRL-CDM indicates a continuous sheath-like surface of interconnected fibers. The white arrow for the shCOL6 points to fragmented and singularized fibers. **(G)** Schematic illustration of AFM (left). Created in BioRender. Schell, C. (2026) https://BioRender.com/x5sqb3y; Violin plot shows Young’s modulus (log_2_ scale) of shCTRL-and shCOL6-CDMs (dots indicate individual measurements of 3 independent experiments, Mann-WhitneyLULtest). **(H)** Representative IF image stained with FN (green) and COL6 (violet), as well as color-coded FN fiber direction of shCOL6 CDM after 24h supplementation of COL6. **(I)** Density plot (left) depicting quantification of FN fiber orientation of CDMs supplemented with COL6 (violet and brown) and COL1 (blue and red). Dot plot (left) of mean AUC within ±30° of centralized peak (N=3 independent experiments, one-way ANOVA with Tukey post-hoc test for multiple comparisons). Bars indicate mean and S.E.M in dot plots or median and quartiles in violin plots; _∗_ – p < 0.05, _∗∗_ – p < 0.01, _∗∗∗_ – p < 0.001 and n.s – not significant.

### ECM composition of fibroblast CDMs is dependent on COL6

For a granular description of ECM composition in CDMs, we followed a similar approach as used in the structural analysis of deposited ECM, but further extended this by additional MS analysis (Figure 4A). Proteomic analysis demonstrated a distinct regulation of various proteins of the core matrisome as well as matrisome-associated proteins. As expected, the three major COL6 chains were significantly depleted in respective knockdown conditions (Figure 4B and Supplementary Figure S4). GO enrichment analysis of regulated ECM proteins indicated a matrix pattern supportive of cellular growth, cell migration (wound healing, chemotaxis) and angiogenesis in control conditions (Supplementary Figure S4). In contrast, depletion of COL6 translated into increased deposition of structural ECM-proteins (e.g., COL5, COL12), ECM-crosslinking enzymes (e.g., LOXL3, LOXL4), as well as proteoglycans (GPC2, GPC4, LUM, BGN – Figure 4B). Aside from these mainly structural ECM-components, also matricellular proteins such as POST or EDIL3 were detected, implicating alterations in signaling properties of respective CDMs. In further validation experiments, we observed an anisotropic deposition pattern for fibrillar collagens (e.g., COL1 and COL5) as well as BGN, as detected for FN in conditions of *COL6* knockdown (Figure 4C-G and Supplementary Figure S4). These findings are most likely related to the inherent mode of assembly and association of respective proteins, which all require tension or mechanical load. On the contrary, tested proteins such as COL4, LAMC1 and FBLN1 were predominantly organized in isotropic patterns (Figure 4E, 4F and Supplementary Figure S4) and showed decreased levels of deposition. These findings indicate that COL6 is essentially involved in the isotropic organization of deposited ECM. Given the striking differences between wild type and COL6-depleted CDM, we further aimed to elucidate potential functional implications via cell-ECM interactions and the consecutive impact on cellular behavior.

**Figure 4.**
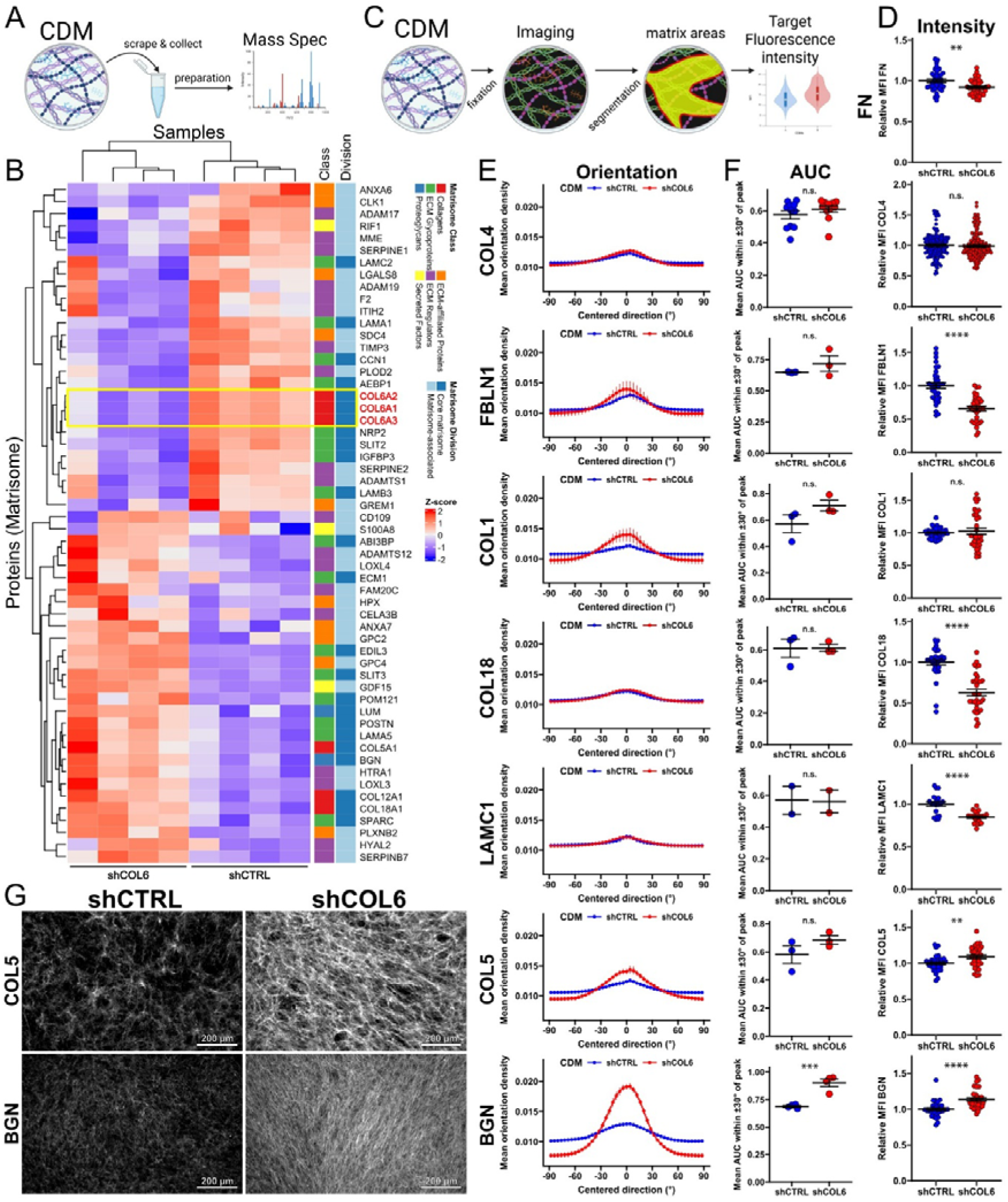
ECM composition of fibroblast CDMs is dependent on COL6. **(A)** Schematic depicting CDM processing for proteomic analysis. Created in BioRender. Schell, C. (2026) https://BioRender.com/x5sqb3y. **(B)** Heatmap presentation of CDM MS-analysis (matrisome-categories are color-coded, COL6A1, COL6A2 and COL6A3 chains are highlighted in red and within the yellow box). **(C)** Schematic depiction of CDM target intensity analysis. Created in BioRender. Schell, C. (2026) https://BioRender.com/x5sqb3y. **(D)** Analysis of relative MFI of respective ECM proteins (image data were pooled from independent experiments, N=11 COL4, N=2 LAMC1, N=4 BGN and N=3 for all else, unpaired t-test with Welch correction for COL5, LAMC1, Mann-Whitney U test for BGN, COL1, COL4, COL18). **(E)** Density plots depicting quantification (mean and S.E.M.) of respective target structure orientation distribution in shCTRL (blue) and shCOL6 (red), calculated from IF images of respective targets (N=11 COL4, N=2 LAMC1, N=4 BGN and for all else N=3 independent experiments). **(F)** Dot plot with quantification of the AUC within ±30° of centralized peak from density curves in (N=11 COL4, N=2 LAMC1, N=4 BGN and for all else N=3 independent experiments, unpaired t-test with Welch correction for BGN, COL5, FBLN1, LAMC1, Mann-Whitney U test for COL1, COL4, COL18). **(G)** IF images of shCTRL-and shCOL6-CDMs stained for COL5 (top) and BGN (bottom). Bars indicate mean and S.E.M in dot plots; _∗∗_ – p < 0.01, _∗∗∗_ – p < 0.001, _∗∗∗∗_ – p < 0.0001 and non-significant (n. s.).

### COL6 propagates the proliferative behavior of cancer cells

To test a potential reciprocal effect of altered CDM composition and structure, we performed reseeding assays to evaluate parameters such as cell morphology, growth and transcriptional changes (Figure 5A). RNA sequencing after cell reseeding of renal cancer cells on respective CDMs showed a pronounced differential transcriptional regulation (Figure 5B). Cells seeded on COL6-containing ECM exhibited enrichment of immunomodulatory processes, including chemokine and cytokine activity, as well as response to hypoxia, as revealed by GO-term analysis (Figure 5C)^26^. Interestingly, most significantly upregulated transcripts in COL6-containing conditions included CXCL1 and CXCL2 genes, which previously have been implicated in tumor-promoting conditions in ccRCC via attracting neutrophil cell populations (Figure 5D)^27^. In addition, factors such as TNFSF14 and LCN2 have also been shown to exert tumor-permissive actions and correlate with aggressive tumor biology^28, 29^. Consistent with these transcriptomic differences, tumor cells showed distinct morphological features upon reseeding. On COL6-containing CDMs, cells formed a more interconnected network structure, whereas on COL6-depleted CDMs, cells aligned with the underlying ECM pattern and adopted a spindle-shaped morphology (Figure 5E-H). Given the significant impact on transcriptional as well as morphological parameters of reseeded tumor cells, we tested the functional relevance of COL6-containing CDMs. And in fact, the proliferative capacity of tumor cells on COL6-depleted CDMs was significantly impaired when compared to COL6-containing conditions (Figure 5I-K). To further validate whether our in vitro observations also apply to the more complex in situ situation, we analyzed the proximity of proliferating tumor cells towards COL6 deposits within ccRCC tissue. Strikingly, the distance of proliferating tumor cells was significantly reduced compared to non-proliferating tumor cells (Figure 5L&M). Together, these observations from in vitro and in situ analysis imply that the pro-proliferative effect of COL6 most likely relates to its modifying function on structural and compositional features of the ECM. This is furthermore supported by a pro-proliferative ECM composition (GO enrichment) and enrichment of SERPINE1 and SERPINE2 as exemplary proteins derived from our CDM proteomics approach (Figure 4B and Supplementary Figure S4)^30^. Moreover, cross-correlation analysis indicated a positive correlation between COL6 chains and respective serpin proteins with classical proliferation markers (e.g. MKI67, CCNB2) in ccRCC (Figure 5N and Supplementary Figure S5). As correlation analysis with TCGA and CPTAC datasets does not contain any spatial information, we transferred our approach using spatial transcriptomics on ccRCC samples and reproduced the positive correlation between COL6, proliferative marker genes and members of the serpin E protein family (Figure 5O&P). In summary, our findings establish COL6 as a central regulator of CDM/ECM functionality with a significant impact on the proliferation of ccRCC tumor cells in vitro and in situ.

**Figure 5.**
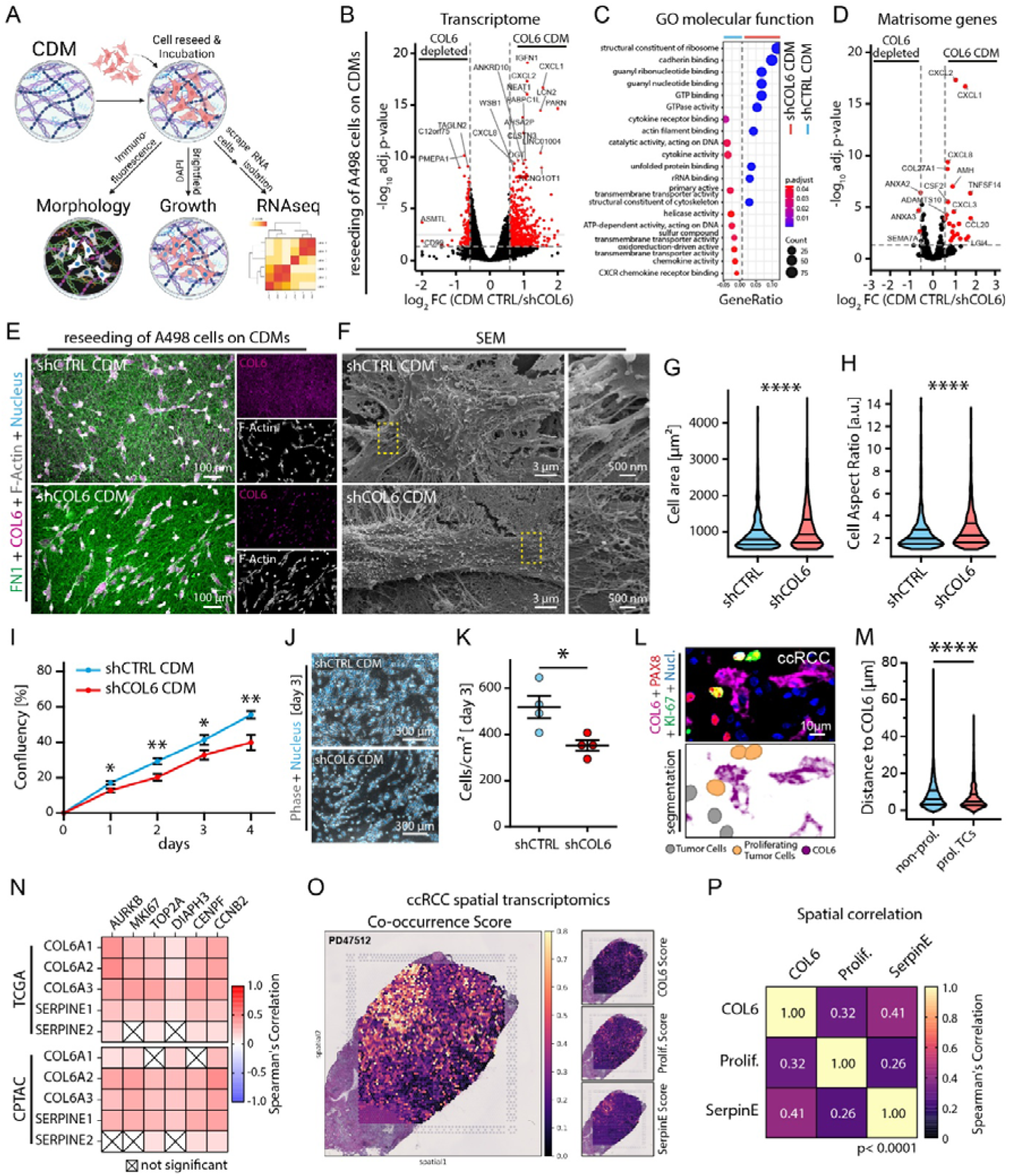
COL6 propagates the proliferative behavior of cancer cells. **(A)** Schematic depicting cancer cell reseeding on TK173-synthesized CDMs and further functional workup (Created in BioRender. Schell, C. (2026) https://BioRender.com/x5sqb3y). **(B)** Volcano plot of RNA sequencing analysis depicting differentially expressed genes of A498 cancer cells after reseeding on TK173-synthesized CDMs (COL6-containing CDM or depleted CDM; red dots indicate significantly regulated genes with adjusted p<0.05 and log_2_ fold change (FC) > |0.5|). **(C)** Gene ontology (GO) gene set enrichment analysis of GO-terms of the molecular function category. **(D)** Volcano plot analysis for filtered matrisome genes. **(E)** IF staining for Nucleus (blue), F-actin (white), FN (green) and COL6 (violet) of A498 cells reseeded on shCTRL- and shCOL6-CDMs. **(F)** Scanning electron microscopy of A498 cells interacting with CDMs (yellow dotted boxes highlight areas of zoom in). **(G&H)** Morphometric analysis of reseeded cells (cell area and perimeter; Violin plots of individual cells of N=3 independent experiments, Mann-WhitneyLULtest). **(I)** Quantification of confluence of A498 cells growing on shCTRL- and shCOL6-CDMs (N=4 independent experiments, unpaired t test). **(J&K)** Representative phase-contrast images with DNA staining (blue) of A498 cells after cultivation for 3 days. Dot plot depicting quantification of A498 cells per cm² grown on CDMs (N=4 independent experiments, unpaired t test). **(L&M)** Representative IF staining of ccRCC patient samples for DNA (blue), PAX8 (red), KI-67 (green) and COL6 (violet) was used to classify and mask (bottom image) for proliferating tumor cells (orange), non-proliferative tumor cells (grey), as well as COL6 (violet). Violin plot depicting quantification of the distance of non-proliferative and proliferating tumor cells to COL6 (Mann-WhitneyLULtest). **(N)** Heatmap depicting Spearman’s correlation analysis of indicated proliferative cell state marker genes with *COL6*, *SERPINE1* and *SERPINE2* genes in the ccRCC TCGA and CPTAC cohorts. **(O&P)** Spatial transcriptomics analysis of ccRCCs for spatial co-occurrence of *COL6*, *SERPINE1*, *SERPINE2* and proliferative cell state marker genes. **(O)** Exemplary spatial mapping of the co-occurrence score on one sample (PD47512) and spatial mapping for the COL6, SERPINE and proliferative gene scores. **(P)** Correlation matrix of the spatial correlation between the COL6, SERPINE and proliferative gene sets (N=5 patients, Spearman’s correlation). Bars indicate mean and S.E.M in dot plots or median and quartiles in violin plots; _∗_ – p < 0.05, _∗∗_ – p < 0.01 and _∗∗∗∗_ – p < 0.0001.

### COL6-rich stromal septa shape the distribution and functional state of tumor-infiltrating T cells in ccRCC

The majority of immune cells enter the tumor microenvironment via intratumoral vessels. Given the pronounced intratumoral vascularization of ccRCC, tumor-infiltrating lymphocytes (TILs) extravasate within the small stromal septa, where pericytes, fibroblasts and immune cells are densely packed^31^. This architectural organization inherently places infiltrating immune cells in close spatial proximity to stromal ECM components such as COL6. To characterize the spatial relationship between major immune cell subsets and COL6 deposition within ccRCC, we employed a multiplexed immunofluorescence (17-plex mIF) approach analyzing 15 cores from 7 patients (Figure 6A-C). At the global level, cell densities of CD3+CD4+, CD3+CD8+, and CD3+CD8+PD1+ T cells positively correlated with COL6 abundance, but overall densities of infiltrating immune cell population were not significantly correlated with COL6 (Figure 6D). Given the spatially restricted deposition pattern of COL6 (Figure 1G), the tumor tissue was subdivided into a COL6-positive stromal compartment (COL6+ area), a 10 µm peristromal boundary zone (COL6 boundary) and the remaining cancer cell compartment (cancer) to enable region-specific analysis. This granular analysis revealed a distinct distribution pattern of the CD3+ T cells, with the highest fraction and density observed in COL6-positive regions, whereas significantly fewer CD3+ T cells were detected intratumorally (Figure 6 E&F). A similar distribution pattern was observed for the CD3+CD8+ T cells, demonstrating a significant correlation of COL6 expression with the spatial distribution of CD3+CD8+ T cells (Figure G&H). In addition, a significantly higher density of CD8+PD1+ T cells was detected in the COL6-positive regions and the adjacent boundary zone compared to the central tumor compartment. Consistently, the fraction of PD1+ cells among the CD8+ T cell population was significantly increased in these areas, with peak aggregation of CD8+PD1+ cells observed within the boundary zone (Figure 6 I-K and Supplementary Figure S6). These findings indicate a spatial accumulation of activated PD1-expressing CD8+ T cells in COL6-rich stromal niches. To functionally interrogate the impact of COL6 on T cell behavior, we employed our established CDM approach and assessed cellular responses ex vivo. PBMCs seeded on CDMs showed invasive behavior after 6 hours (Figure 6L). In analogy, TILs (tumor infiltrating T-cells) exhibited pronounced filopodial protrusion formation, as well as larger cell area and perimeter on COL6-depleted CDMs after 4 hours (Figure 6M&N and Supplementary Figure S6). Notably, transmigration assays with peripheral T cells demonstrated that COL6 within the ECM supported rather than restricted cell migration (Figure 6O and Supplementary Figure S6). Taken together, these findings argue against a direct COL6-mediated trapping mechanism and instead suggest that COL6 contact modulates the functional state of T cells and thereby inhibiting further infiltration into the cancer cell compartment. Finally, analysis of immune checkpoint inhibitor (ICI)-treated ccRCC samples indicated infiltration of CD3+CD8+PD1+ and CD3+CD4+ T cells into the cancer compartment, associated with cancer necrosis (Figure 6P and Supplementary Figure S6). However, ICI-treatment is also associated with elevated COL6 deposition within most tumor areas, which was particularly prominent in regions of residual fibrosis where viable tumor cells were absent. This observation is confirmed by analysis of COL6 deposition in vital tumor tissue of 3 ICI-treated patients (Figure 6Q). Overall, these observations point to an intricate interplay between stromal COL6 and T cells, and further suggest that ICI treatment regimens directly remodel COL6-associated features of the TIME.

**Figure 6.**
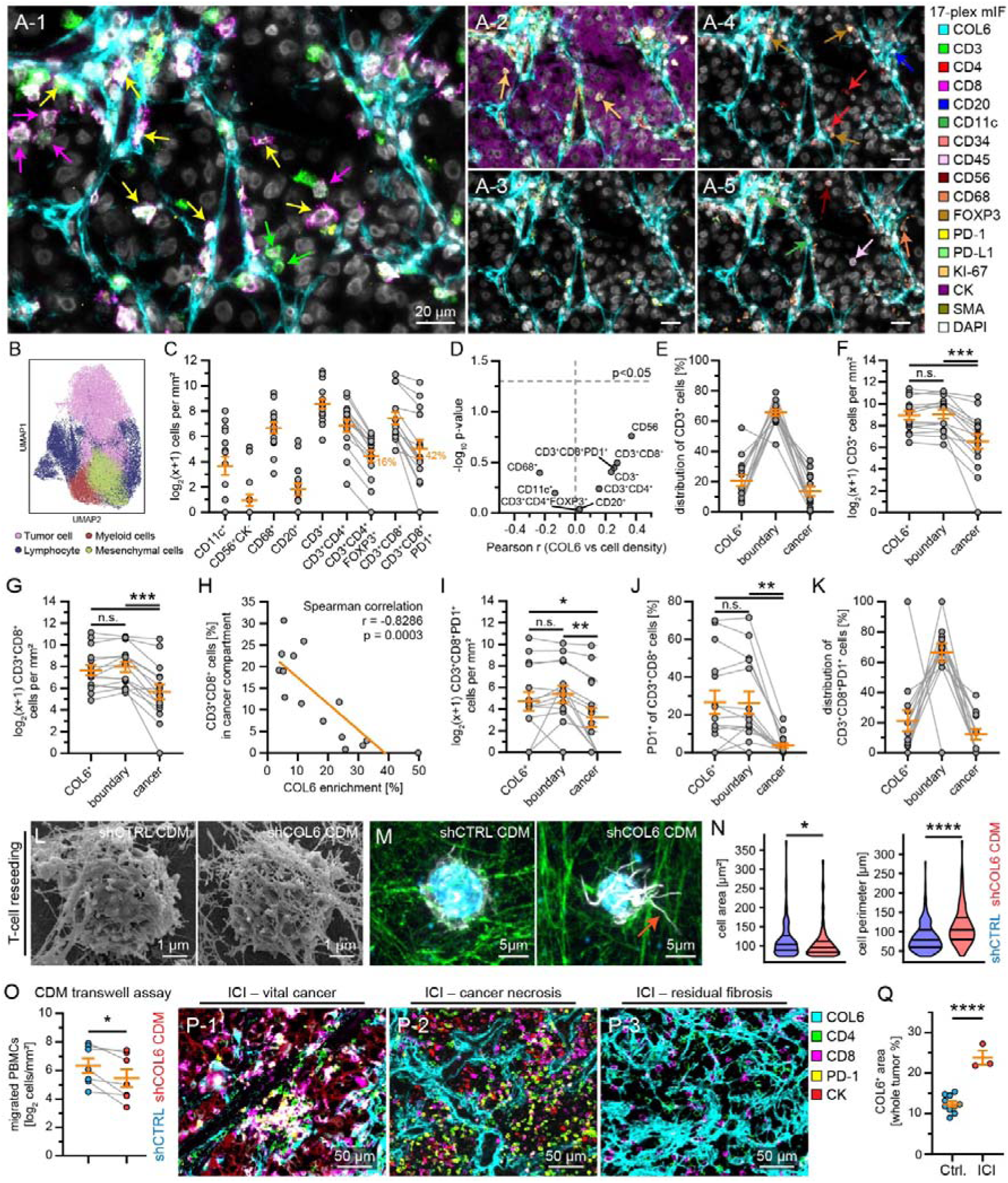
COL6-rich stromal septa shape the distribution and functional state of tumor-infiltrating T cells in ccRCC. **(A)** 17-plex SeqIF analysis for COL6 and the spatial distribution of immune cell populations in ccRCC was applied on 15 cores from 7 cancer cases. Representative images show IF stainings of indicated cell and compartment markers. Arrows indicate representative cell types: light green – CD3+CD8- (A-1), purple – CD3+CD8+ (A-1), yellow – CD3+CD8+PD1+ (A-1), light orange – KI67+ (A-2), red – CD4+ (A-4), brown – CD4+FOXP3+ (A-4), dark blue – CD20+ (A-4), dark green – PDL-1+ (A-5), dark red – CD56+ (A-5), orange – CD68+ (A-4). **(B)** UMAP of major segmented cell populations (53 102 cells, tumor cells – pink, myeloid cells – red, lymphocytes – blue and mesenchymal cells – green). **(C)** Log_2_ density of indicated immune cell population in analyzed samples (dots indicate N=15 cores in graphs C- K). Percent numbers indicate the mean density of CD3+CD4+FOXP3+ cells of total CD3+CD4+ cells and CD3+CD8+PD1+ cells of total CD3+CD8+ cells. **(D)** Correlation analysis of whole tumor immune cell densities with COL6 expression (Pearson correlation). **(E)** Relative distribution of CD3+ cells to the COL6+ stromal compartment, peristromal boundary zone (COL6 boundary) and the remaining cancer cell compartment (cancer) in ccRCC. **(F**) Absolute density of CD3+ cells in indicated compartments (N=15, RM one-way ANOVA with Geisser-Greenhouse correction and Tukey’s multiple comparison test is used in graphs F, G, I and J). **(G)** Absolute density of CD3+CD8+ cells in the indicated compartments. **(H)** Correlation analysis of absolute CD3+CD8+ cell density in the cancer compartment with tumor COL6 expression (Spearman correlation). **(I)** Absolute density of CD3+CD8+PD1+ cells in the indicated compartments. **(J)** Percentage of CD3+CD8+PD1+ cells of total CD3+CD8+ cells in indicated compartments. **(K)** Relative distribution of CD3+CD8+PD1+ cells to indicated compartments. **(L)** Scanning electron microscopy (SEM) images of peripheral blood mononuclear cells (PBMCs) seeded on shCTRL- and shCOL6-CDMs after incubation for 6 hours. **(M)** IF images of T cells incubated on shCTRL- and shCOL6-CDMs stained for DNA (blue), FN (green), and F-actin (white). T cells incubated on shCOL6-CDMs exhibit an increased formation of protrusions (white arrow). **(N)** Quantification of T cell morphology based on cell area and cell perimeter, after incubation with respective CDMs (Violin plots of analyzed cells of N=4 independent experiments, Mann-WhitneyLULtest). **(O)** Quantification of transwell migration of PBMCs through CDMs synthesized by shCOL6 and shCTRL fibroblasts (N=7 replicates of 4 independent experiments, paired t-test). **(P)** SeqIF analysis of an immune checkpoint inhibitor (ICI – Nivolumab & Ipilimumab) treated ccRCC patient sample. Representative images of regions with vital cancer, immune infiltration and cancer necrosis, and residual fibrosis are shown (markers as indicated). **(Q)** Quantification of COL6 positive area in the vital cancer compartment of whole tumor section of 3 ICI-treated patients and 10 control ccRCCs (dots indicate individual patients analyzed, Mann-WhitneyLULtest). Bars indicate mean and S.E.M in dot plots or median and quartiles in violin plots; _∗_ – p < 0.05, _∗∗_ – p < 0.01, _∗∗∗_ – p < 0.001, _∗∗∗∗_ – p < 0.0001 and non-significant (n. s.).

### Relevance of COL6 in the context of conventional treatment regimens of ccRCC

Current therapeutic approaches for advanced ccRCC encompass tyrosine kinase inhibitors and/or combinatory immune checkpoint therapy^32, 33, 34^. Survival analysis of nivolumab-treated patients revealed that high COL6 expression (in particular COL6A1 and COL6A2) correlated with impaired outcome, whereas this association was not observed in patients receiving mTOR inhibitor therapy (Figure 7A). Remarkably, high COL6 expression levels were associated with a progressive disease course in our proteomics analysis (Supplementary Figure S7). These data prompted us to investigate whether established ccRCC treatment regimens directly modulate the stromal COL6 compartment. In a single case of neoadjuvant TKI treatment (primary tumor), markedly reduced COL6 levels were apparent in the resected tumor tissue (Supplementary Figure S7). This observation is in marked contrast to the elevated COL6 deposition levels found in ICI treatment regimens (Figure 6Q). To mechanistically dissect the effect of TKI treatment on stromal fibroblasts, we treated our TK173 fibroblast model with cabozantinib and assessed resulting alterations in CDM composition. Cabozantinib treatment led to a pronounced depletion of COL6 within the CDM relative to control conditions. This effect was validated using sunitinib, demonstrating a consistent class effect for TKIs, whereas the mTOR inhibitor temsirolimus exerted a minor suppressive effect on COL6 deposition (Figure 7B-D and Supplementary Figure S7). To delineate the underlying transcriptional mechanisms, we performed integrative transcriptomic profiling of cabozantinib-treated TK173 fibroblasts and ex vivo ccRCC tumor models (Figure 7E-I). Cabozantinib treatment induced a significant downregulation of all three major COL6 chains in fibroblasts, while other collagen subtypes remained largely unaffected or were even increased (Figure 7F&H and Supplementary Figure S7). Remarkably, transcriptomic analysis of cabozantinib-treated ex vivo ccRCC samples closely mirrored these in vitro findings, with reduced expression of all three COL6 chains (Figure 7G&H). Beyond direct ECM remodeling, deconvolution analysis of both primary treated ccRCC samples and TK173 cells demonstrated complex co-regulation of central immunomodulatory mediators, including key chemokine ligands (e.g., CX3CL1, CXCL10, VEGFA) and receptors (e.g., ICOS, LAG3, CD27 – Figure 7I). The convergent regulation of CX3CL1 across both primary tumor tissue and fibroblast cell lines is particularly noteworthy, as it suggests that TKI-mediated stromal remodeling may directly reshape immune cell recruitment and the overall immune microenvironment of ccRCC. Reduced COL6 deposition in cabozantinib-treated ccRCC tissue was further confirmed at the tissue level (Figure 7J&K). Taken together, given that elevated COL6 expression associates with poor patient survival and promotes tumor cell proliferation, our finding that cabozantinib selectively suppresses COL6 expression in stromal fibroblasts reveals a previously underappreciated mechanism of TKI action. Our findings imply that TKI treatment exerts functions beyond direct tumor cell targeting to encompass functionally relevant remodeling of the matrisomal and immune compartments of the ccRCC TIME (Figure 7L).

**Figure 7.**
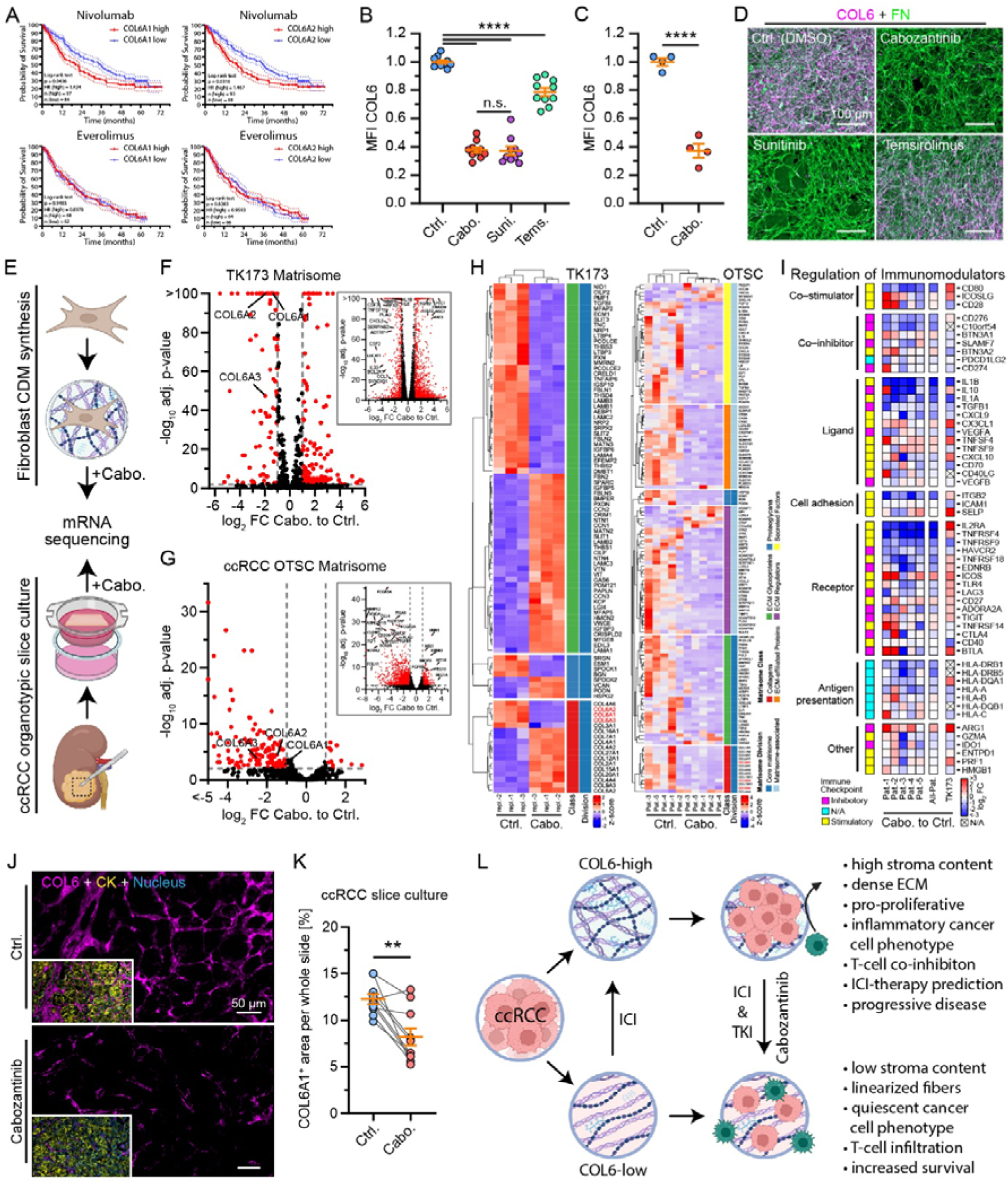
Relevance of COL6 in the context of conventional treatment regimens of ccRCC. (**A**) Survival analysis for high or low expression of *COL6A1* and *COL6A2* in the CheckMate 025 study cohort of Nivolumab (anti-PD-1) or everolimus (mTOR inhibitor) treated patients^64^. Graphs show Kaplan-Meier plots and Log-rank (Mantel-Cox) tests. **(B-D)** Analysis of COL6 deposition in CDMs treated with cabozantinib (Cabo.), sunitinib (Suni.), temsirolimus (Temsi.), or DMSO (Ctrl.) treated TK173 cells. (B) Dot plot depicting relative median fluorescence intensities (MFI) of COL6 in CDMs of respectively treated cells in the screening experiment (N=10 regions of interest, one-way ANOVA with Tukey’s multiple comparison test). (C) Dot plot depicting mean relative MFI of COL6 in CDMs of respectively treated TK173 cells (dots indicate mean of N=4 independent experiments, unpaired t test). (D) IF images stained for FN (green) and COL6 (violet) of synthesized CDMs of TK173 cells treated with the indicated drugs. **(E)** Schematic description of RNA sequencing analysis of cabozantinib or DMSO control-treated TK173 fibroblasts and acute organotypic slice cultures (OTSCs) of ccRCCs (Created in BioRender. Schell, C. (2026) https://BioRender.com/x5sqb3y). **(F&G)** Volcano plot of differential gene expression analysis of matrisome genes in cabozantinib-treated TK173 cells and ccRCC OTSCs. The inserts depict a volcano plot of all genes. Red dots indicate significantly regulated genes with adjusted p<0.01 and log_2_ fold change (FC) > |1|). **(H)** Heatmap analysis of significantly regulated matrisome genes in cabozantinib-treated TK173 cells and ccRCC OTSCs (genes with adjusted p<0.0001 and fold change > |2| are depicted for TK173 cells or with adjusted p<0.01 and fold change > |1.5| for OTSCs). **(I)** Heatmap analysis of log_2_ expression fold changes (FC) of immunomodulatory genes^75^ in TK173 cells and ccRCC OTSCs (N/A – not announced). **(J&K)** IF validation of COL6 regulation in cabozantinib-treated ccRCC OTSCs. Images show IF staining for Nuclei by Hoechst (blue), CK (yellow) and COL6A1 (violet) of OTSCs treated with the indicated drugs. Quantification of COL6A1-positive areas in OTSCs of 10 ccRCC patients (dots indicate individual patients analyzed, ratio paired t test). **(L)** Schematic summary illustrating the findings of the study, highlighting COL6 expression in tumor stroma as well as the remodeling of the ECM architecture in dependence of COL6 abundance and the subsequent implications for cancer and immune cells (Created in BioRender. Schell, C. (2026) https://BioRender.com/x5sqb3y).

## Discussion

The role of the extracellular matrix in the context of cancer biology is mainly based on observations made in stroma-rich tumors, which are characterized by a dense accumulation of ECM and overall stiff rigidity^35^. Interestingly, also rather stroma-poor tumors exhibit distinct ECM signatures correlating with tumor aggressiveness, indicating matrisome functions beyond sole mechano-signaling. Clear cell renal cell carcinoma is a prime example of a stroma-poor tumor, while previous studies clearly demonstrated the relevance of ECM alterations for parameters such as metastasis and patient outcome^13, 36^. Here, we aimed to delineate the implications of ECM composition in ccRCC and focused on the tumor-instructive role of COL6.

Employing bulk proteomics on a ccRCC patient cohort, we demonstrate that COL6 is significantly accumulating in the tumor tissue (Figure 1 and Supplementary Figure S1). While previous studies have documented COL6 upregulation and its impact on tumor cell biology^5, 37, 38, 39^, a critical distinction must be drawn regarding its cellular origin. By combining available scRNAseq datasets with multiple imaging modalities, our data consistently indicate that the predominant source of COL6 is detectable in the interstitial compartment of ccRCC tumor tissue (Figure 1 and Supplementary Figure S1). This spatial context is of substantial importance as prior work has predominantly focused on tumor cell-intrinsic roles of COL6, underestimating the functional significance of stromal COL6 deposition. To delineate intrinsic and extrinsic signaling effects of COL6, we employed genetic titration strategies and consistently observed that COL6 knockdown primarily translates into altered growth behavior accompanied by transcriptional alterations affecting biological functions such as focal adhesions or collagen formation (Figure 2).

Given the predominant deposition of COL6 within the tumor stroma of ccRCC, a major aspect of its function likely lies in the structural organization of the ECM. This effect of COL6 is known in the context of its physiological functions via organizing the three-dimensional tissue architecture of skeletal muscles, tendons, bone and cartilage^16^. Moreover, recent work has demonstrated drastic ECM-alterations in COL6-related muscular dystrophies^40, 41^. However, to the best of our knowledge, how stromal COL6 specifically influences tumor cell biology in ccRCC has remained poorly understood.

We found that COL6 is essentially required to maintain the structural complexity of the FN-network within CDMs (Figure 3). These observations are in line with previous reports on CDM models derived from COL6-related muscular dystrophies (COL6-RD) patients, implying a conserved biological mechanism^41^. In addition, we demonstrate that the matrix modulating capabilities of COL6 are directly mediated via the protein itself and its deposition in the ECM, since supplementation of COL6-depleted CDMs with COL6 restored the ECM structural complexity (Figure 3). These effects may partly rely on the capacity for self-assembly shared by certain ECM proteins, particularly collagens^42^. The pronounced structural alterations of CDMs upon COL6 depletion also implied changes within the composition of the respective ECM. Here, we employed a combined approach using proteomics of CDMs and immunofluorescence analysis (Figure 4). Our data show that COL6 has a major instructive role on the composition of deposited ECM, which also altered reciprocal cell-matrix interactions. Interestingly, testing the growth behavior of cancer cells on COL6-depleted ECMs translated into altered cellular proliferation rates, while parameters such as matrix stiffness or related mechano-signaling were not affected (Figure 4&5). Many of the regulated proteins are part of the core matrisome, such as collagens, laminins and proteoglycans. Furthermore, ECM regulators (e.g., serpins and lysyl oxidase-like (LOXL) proteins) were differentially detected, indicating a major impact on structural remodeling. Notable regulated proteins include insulin-like growth factor binding protein 3 (IGFBP3), which exhibits both growth-supportive^43^ and inhibitory functions^44^. In addition, members of the serpin E protein family (SERPINE1, SERPINE2) were not only co-regulated with COL6 in our CDM approaches, but also spatially co-localized within ccRCC tissue and corresponded to a proliferative gene signature in spatial transcriptomic data sets (Figure 4&5). This is in line with previous work demonstrating that levels of SERPINE2 are indicative of aggressive tumor biology in advanced renal carcinoma^30^.

Most immune cells enter the tumor microenvironment via intratumoral vessels, and given the pronounced intratumoral vascularization of ccRCC, tumor-infiltrating lymphocytes (TILs) extravasate within the small stromal septa^31^. This architectural organization inherently places infiltrating immune cells in close spatial proximity to stromal ECM components such as COL6. Granular spatial analysis of ccRCC tumor tissue demonstrated that the majority of CD3+ T cells localized near COL6-rich stromal septa, whereas significantly fewer T cells were observed within the center of tumor acini (Figure 6). This distributional pattern was similarly observed for CD3+CD8+ and CD3+CD8+PD1+ T cells, suggesting preferential T cell clustering at the boundary zone with concurrent features of chronic activation and exhaustion. Conversely, only a minor fraction of PD1+ effector T cells was identified within the tumor center zone in direct contact with malignant cells. These findings are consistent with previous studies showing that COL6-modified matrix induces CD8+ T cell dysfunction in sarcoma models^45^ and that COL6 correlates with reduced T cell motility^46^. To functionally interrogate the impact of COL6 on T cell behavior, we employed our established CDM model and assessed primary T cell responses ex vivo. T cells seeded on COL6-depleted CDMs exhibited pronounced filopodial protrusion formation (Figure 6), a morphological phenotype indicative of enhanced environmental engagement and potentially facilitating formation of the immunological synapse^47^. Notably, transmigration assays demonstrated that COL6 within the ECM supported rather than restricted cell migration (Figure 6). This finding argues against a simplistic COL6-mediated physical trapping mechanism and instead indicates that COL6 contact modulates functional T cell states, potentially promoting an exhausted phenotype rather than imposing a mechanical barrier. Such a regulatory event may be transmitted through LAIR1, an immunosuppressive receptor upregulated by interaction of T cells with collagens in cancer, potentially conferring resistance to ICI therapy^48, 49^. Taken together, these findings reveal an intricate interplay between stromal COL6 and infiltrating T cells within the ccRCC TME, in which COL6-rich septa shape both the spatial distribution and functional state of T cells. Furthermore, ICI treatment regimens appear to directly remodel COL6-associated features of the TME (Figure 6), with potential implications for immune cell re-engagement within treated tumors.

The standard treatment for advanced ccRCC often involves immune checkpoint inhibitors, such as ipilimumab (CTLA-4 blockade), nivolumab (PD-1 blockade), or a combination of both, or TKIs, such as cabozantinib^32^. Cabozantinib inhibits vascular endothelial growth factor receptor 2 (VEGFR2), MET and AXL^50^, thereby reducing tumor growth, metastasis and angiogenesis^51, 52^. Here, we present further evidence of how cabozantinib may influence tumor growth and immunity. We show that cabozantinib treatment of renal fibroblasts results in impaired ECM deposition, characterized by reduced COL6 expression. Furthermore, the ECM architecture of cabozantinib treated fibroblasts resembled the loss of FN fiber complexity observed in COL6 depletion (Figure 7). Critically, this COL6-suppressive effect of cabozantinib was not restricted to in vitro fibroblast cultures. Examination of ex vivo treated tumor samples demonstrated that cabozantinib treatment reduces COL6 expression in a more physiologically relevant context (Figure 7), further supporting the translational relevance of these findings. Given that COL6 depletion in CDMs influences cancer cell growth and T cell interaction (Figures 5&6), the reduction of COL6 upon TKI treatment may represent a previously unrecognized, clinically relevant mode of action of cabozantinib, potentially involving ECM remodeling and modulation of the immune microenvironment (Figure 7). Together, our observations indicate the essential relevance of spatial and compositional ECM dynamics in tumor progression with significant implications for improving therapeutic strategies in ccRCC.

## Methods

### Human samples

The study was conducted according to the guidelines of the Declaration of Helsinki and approved by the Institutional Ethics Committee of the University Medical Center Freiburg (EK 18/512, EK 21/2188). Informed consent was obtained from all subjects involved in the study.

### Histological analysis of ccRCC samples

Formalin-fixed, paraffin-embedded (FFPE) ccRCC tissue samples that were included in the proteomics cohort were first stained with hematoxylin and eosin (H&E). Only cases with both high-quality H&E sections and corresponding proteome data were selected for histological image analysis (n=23). Histological image analysis followed the workflow previously developed and published^11^, including manual tumor annotation, pixel-based tissue classification in QuPath^53^, StarDist-based cell segmentation and supervised object classification, which was applied accordingly to the samples of the ccRCC cohort used in this study.

### Antibodies and dyes

Antibodies used in this study are described in the respective method sections. Further detailed information is provided in Supplementary Spreadsheet S1.

### Immunofluorescence

Immunofluorescence (IF) staining was performed on cells, CDMs, reseeded cells, and FFPE tissue sections. Cells and CDMs were fixed in 4% paraformaldehyde (PFA, 15714-S, Electron Microscopy Sciences) in phosphate-buffered saline (PBS) with 1 mM CaCl_2_/MgCl_2_ for 20 min, and permeabilized with 0,1% Tergitol (A9780, AppliChem). FFPE tissue sections were deparaffinized before staining. HIAR was done where required using citrate buffer (pH 6) or Tris-EDTA buffer (pH 9) in a pressure cooker. For LAMC1 staining of CDMs HIAR was conducted in Tris-EDTA at 95°C for 30 min in a steamer. Samples were blocked with 5% bovine serum albumin (BSA) (11926.03, Serva) in PBS for 1 h and incubated overnight at 4°C with primary antibodies diluted in 5% BSA in PBS. Following washes with PBS, fluorophore-conjugated secondary antibodies (Alexa Fluor dyes, Thermo Fisher Scientific) were applied for 1h. Nuclei were stained with Hoechst 3342 (H3569, Thermo Fisher Scientific) and, where indicated, F-actin with phalloidin (Thermo Fisher Scientific). Imaging was performed using an Axio Observer microscope (Carl Zeiss AG) equipped with an Apotome 2 device for wide-field optical sectioning.

### Multiplex Immunofluorescence

Cyclic immunohistochemistry (cycIHC) and OPAL 6-Plex staining (NEL811001KT, Akoya Biosciences) were performed as previously described^7, 54^. Briefly, FFPE tissue sections were cut, mounted on slides, deparaffinized in xylene, and rehydrated. Iterative staining was repeated for multiple markers on the same section (Supplementary Spreadsheet S1). Whole-slide imaging was conducted and image analysis was done with QuPath 0.4.4. The positive COL6 area was quantified using channel thresholding. For proximity analysis, cell segmentation was performed and cell classification based on PAX8 and Ki-67 was done. PAX8 was used to classify tumor cells, and proliferating tumor cells were defined as PAX8^+^Ki-67^+^. Distances between tumor cells and COL6-positive regions were calculated using the **“**Signed distance to annotations 2D” tool. Cells within the COL6-positive area were assigned a distance of 0 (instead of native negative values) for visualization purposes. Sequential immunofluorescence (SeqIF) was performed using the COMET platform (Lunaphore, Biotechne) on FFPE tissue sections derived from a previously published ccRCC TMA cohort and one exemplary neoadjuvant immune checkpoint inhibitor-treated ccRCC patient^7^. Staining was conducted using the SPYRE Antibody Core Panel (Lunaphore) complemented with additional primary antibodies validated in our lab to enable up to 17-plex imaging (Supplementary Spreadsheet S1). Sequential staining cycles were performed according to the manufacturer’s protocol. Raw image data were processed using the HORIZON software (Lunaphore), including background subtraction. To account for intratumoral heterogeneity and to preserve spatial resolution, each TMA core was treated as an independent analytical unit. This approach enabled the assessment of COL6-associated spatial features across distinct tumor regions. For image analysis, the tumor tissue was sub-segmented into COL6-positive stroma, a 10 µm peristromal zone surrounding the COL6-positive stroma, and the remaining tumor compartment. Only vital tumor tissue was analyzed, excluding the pseudocapsule or region of fibrosis. Cell segmentation was done using InstaSeg in QuPath v0.6.0. Immune cell populations were classified using marker-based single or combinatorial threshold classifiers. Cell densities, relative frequencies, spatial compartmentalization, and compartment sizes were quantified for downstream analysis.

### Cell Culture

Human renal cancer cell lines A498 (HTB-44, ATCC,) and 786-O (CRL-1932, ATCC) as well as human renal fibroblasts TK173 (RRID: CVCL_C8FA) and human renal proximal tubule epithelial cells (RPTEC/TERT1, ATCC, CRL-4031) were used and obtained as recently described^11^. A498, 786-O, TK173 and RPTEC cell lines were cultured below 80% confluence using culture medium comprising RPMI-1640 (61870036, Thermo Fisher Scientific) supplemented with 10% FCS (S0615, Sigma-Aldrich) and 1% penicillin-streptomycin (P/S, 15140122, Thermo Fisher Scientific). Cell lines comprising *COL6A2*-knockdown (KD) were transduced with small hairpin RNAs in the pLKO.1 lentiviral vector designed by The RNAi Consortium (TRC). Plasmid transduction in order to generate *COL6A2*-KD and overexpressing *COL6A2* (oeCOL6A2) as well as respective controls, was achieved with lentiviral particles produced via PEI-transfection in human embryonic kidney (HEK) 293T cells (CRL-3216, ATCC). HEK293T cells were cultured in DMEM (31966-021, Thermo Fisher Scientific) supplemented with 10% FCS. All cell lines were cultured at 37°C in a 5% CO2 humidified atmosphere. Routine testing of cell lines for mycoplasma contamination was performed using a polymerase chain reaction (PCR)-based detection kit (Mycoplasma PCR detection kit, Hiss Diagnostics GmbH, Germany). Generation of peripheral blood mononuclear cells (PBMC) and Tumor-infiltration Lymphocyte (TIL) isolation and expansion was performed as previously reported^55^.

### Expression Plasmids

Cell lines comprising *COL6A2*-knockdown (KD) were generated by expressing different validated small hairpin RNA (shRNA) targeting *COL6A2* cloned into pLKO.1 lentiviral vector (purchased by Horizon Discovery – The RNA Consortium, clone ID: TK173-shCOL6A2-1: TRCN0000116848 (antisense-sequence: TTTCTCTCGGTAGTGTCCGGC), TK173-shCOL6A2-2: TRCN0000116849 (antisense-sequence: TTGACCACGTTGATGACGAAG), 786-O-shCOL6: TRCN0000116850 (antisense-sequence: ATGAAGCGGTCGAAGAAGCCG). pLKO.1 control (further termed “shCTRL”) was a gift from William Hahn (Addgene plasmid, 42559; RRID: Addgene_42559). oeCOL6A2 cells were generated by expressing human hCOL6A2 (NM_001849.4) in pCSdest. COL6A2_pCSdest was a gift from Roger Reeves (Addgene plasmid, 53903; RRID: Addgene_53903)^56^. COL6A2 and luciferase control (oeCTRL)^57^ were subcloned into pLenti-C-Myc-DDK-P2A-Puro expression vector backbone (PS100092, OriGene Technologies).

### Western Blot

Western blotting was performed as described previously^11^. Briefly, cells were lysed in RIPA buffer, and protein concentrations were determined by BCA assay. Equal protein amounts were separated by SDS-PAGE, transferred to PVDF membranes, and probed with the indicated primary antibodies, followed by HRP-conjugated secondary antibodies. Signals were detected by enhanced chemiluminescence using a digital chemiluminescence imager.

### Generation and preparation of cell-derived matrices

Cell-derived matrices (CDMs) were generated using TK173 and 786-O cell lines as previously described^25^. In brief, glass coverslips, 35 mm, low, glass-bottom dishes (80137, Ibidi) or transwell inserts, 8 µm pore for 24-well plate (662638, Greiner) were coated with gelatin (G1890, Sigma-Aldrich) and crosslinked with GA (glutaraldehyde, 4157.2, Carl Roth) for imaging assays, whereas uncoated 10 cm dishes were used for CDMs used for proteomic or transcriptomic analysis. Cells were seeded to reach confluence within 1–3 days and then cultured for 7 days (if not indicated otherwise) in medium supplemented with 50 μg/ml vitamin C (A4544, Sigma-Aldrich), which was renewed daily. CDMs were decellularized by extraction with 0,5% Tergitol and 20 mM NH_4_OH in PBS, followed by DNase I treatment (15 U/ml in PBS + 1 mM CaCl_2_/MgCl_2_ - A3778, AppliChem) and excessive washing to remove residual DNA and cell detritus, and the resulting matrices were used for downstream analyses. For treatment experiments, TK173 cells were treated during the CDM generation with respective inhibitors or an equal volume of DMSO. Inhibitors used and respective working concentrations are 5 µM cabozantinib (S1119, Selleck Chemicals, Houston, TX, USA), 10 µM sunitinib (S1042, Selleck Chemicals, Houston, TX, USA) and 10 µM temsirolimus (S1044, Selleck Chemicals, Houston, TX, USA), medium was changed every second day. For CDM supplementation experiments, pre-formed CDMs on coverslips were overlaid with neutralized COL1 (5005, Advanced Biomatrix) or COL6 (009-001-108, Rockland Immunochemicals) solutions (250 μg/ml in PBS), incubated for 24 h at 37°C, and subsequently processed for immunofluorescence staining. For the cell reseeding experiment, cells were reseeded in culture medium and incubated at 37°C, 5% CO_2_ in a humidified incubator for the indicated time periods.

### Analysis of cell morphology, focal adhesions, proliferation, and matrix structure

Analysis of both cellular and focal adhesion morphology was conducted by applying previously described protocols and CellProfiler 4.7.2^54^. BrdU cell proliferation (ab126556, abcam, Cambridge, UK) analysis was performed following protocol instructions. Clonogenic capability of cells over 7 days was analyzed by staining with 1% crystal violet and analysis in Fiji ImageJ v1.54f using standard procedures. CDM thickness was assessed by measuring the distances between the top and bottom z-layers of z-stack images. Analysis of CDM fiber orientation was performed using Fiji ImageJ v1.54f and plugins “Directionality” for quantification of orientation histograms and angular dispersion, as well as “OrientationJ” for the generation of color-coded fiber images. To examine the morphology of cells on CDMs, cells were seeded in culture medium and incubated for 4 h (PBMCs for 6 h) before washing once with PBS and processing for IF staining or respective imaging protocol. Morphological analysis was conducted using CellProfiler 4.7.2. To observe cell confluency on CDMs over time, brightfield images were taken daily and analyzed using Fiji ImageJ v1.54f. For growth analysis by cell counting, cells were stained using Hoechst and counted using QuPath 0.5.1. For Transwell migration analysis, CDM was synthesized on both sides of 8 μm pore membrane transwells (662638, Greiner Bio-One) as described above, excluding gelatin coating. Transwell migration was performed and analyzed as previously described, with an adjusted incubation time of 48 h for PBMCs^11^.

### Super resolution imaging of CDMs (IF/SEM)

A ZEISS LSM980 MP AiryScan 2 microscope (Carl Zeiss AG) was used for super-resolution imaging of CDMs (Plan-Apochromat 63x objective, 130 nm/pixel scan). Z-stacks of 170 nm per z-layer were imaged, processed using the Airyscan SR 3D tool and converted to maximum intensity projection. Scanning electron microscopy (SEM) of CDMs was conducted at the EM core facility of the Department of Nephrology (Faculty of Medicine, University of Freiburg, EMcore RI_00555). Generation of CDMs and reseeding of cells was performed as described for IF analysis. Samples for SEM were fixed in 4% PFA and 1% GA in PBS for 24 hours. Fixed samples were further processed for SEM and imaged using a Quattro SEM (Thermo Fisher Scientific) as previously described^57, 58^.

### Atomic Force Microscopy of CDMs

A JPK NanoWizard 3 atomic force microscope (Bruker Nano GmbH) with a soft, rectangular cantilever with a cylindrical tip (5 μm end radius) and a spring constant of *k* = 0.265-0.299 N/m (SAA-SPH-5UM, Bruker) was used to acquire on average 5 – 10 force maps per sample; each force map consisted of a minimum of 16 force-indentation curves (16 – 64) captured over a 10 μm × 10 μm measurement area in an 8 × 2 or 8 x 8 grid (corresponding to 1.25 μm lateral spacing). Measurement sites were selected to characterize as much of the CDM as possible while avoiding regions with cellular debris or where the matrix was not well adhered to the dish or showed voids. Measurement settings used were 1.8 nN of indentation force applied at 1.0 μm/s indentation speed in force mapping mode resulting in an average indentation depth of about 1.5 -2.0 μm; parameters were chosen to avoid inelastic deformation of the matrix. The spring constant of the cantilever was pre-calibrated by the producer (Bruker), the sensitivity was calibrated daily prior to each sample measurement using the JPK calibration tool’s thermal noise method based on the procedure described by Hutter and Bechhoefer^59^. The resulting force curves were analyzed using JPK analysis software to calculate bulk tissue Young’s Modulus by fitting the Hertz/Sneddon model for contact mechanics, taking the indenter shape into account.

### Ex vivo culture and treatment of ccRCC patient samples

Tumor tissue was processed for ex vivo organotypic tissue slice culture (OTSC) as previously described^60, 61^. Briefly, specimens were cut into 0.5 cm^2^ pieces, embedded in 2% agarose, and sectioned at 300 µm thickness using a Leica VT1200 S vibratome (Leica Biosystems). Slices were placed individually onto cell culture inserts (PICM0RG50, Millipore) in 6-well plates and cultured for 5 days in RPMI medium supplemented with 10% FCS and 1% P/S (1 ml per well). Tissue slices were treated with either 0.1% DMSO as a negative control, 25 µM cabozantinib for RNA sequencing analysis or 10 µM cabozantinib for 5 days for IF analysis. Following treatment, slices were fixed in 4% PFA, embedded in paraffin (FFPE), and processed for IF staining as described above. Image analysis was performed using QuPath 0.4.4, and COL6-positive areas were quantified by channel thresholding.

### Gene expression and prognosis analysis

Kaplan-Meier plots of *COL6A1*, *COL6A2* and *COL6A3* mRNA expression in ccRCC were generated using the GEPIA2 tool, which is based on the TCGA KIRC dataset^62^. Kaplan-Meier plots for *SERPINE1* and *SERPINE2* were generated using analyses and best expression cut-offs from the Human Protein Atlas, which is based on the TCGA KIRC dataset^63^. Kaplan-Meier plots of *COL6A1*, *COL6A2* and *COL6A3* expression in the CheckMate 025 study cohort of Nivolumab (anti-PD-1) or everolimus (mTOR inhibitor) treated patients were generated using previously published datasets^64^. Mean expression of *COL6* genes was calculated and used for the separation of tumors into high and low expression of *COL6* genes in respective cohorts. RNA sequencing expression data for COL6, SERPINE and proliferation (cycling cell state) associated genes in the TCGA and CPTAC ccRCC cohort were derived from the cBioPortal platform (CPTAC GDC 2025 and TCGA, Firehose Legacy datasets)^65^. Kaplan-Meier plots, heatmaps, Log-rank (Mantel-Cox) tests and Spearman correlation analyses were performed using GraphPad Prism v8 software.

### Single Cell RNA sequencing analysis

Single-cell RNA sequencing (scRNA-Seq) data were assessed and analyzed via the CELL×GENE platform from a previously published cohort of 7 ccRCCs^66, 67^. Analysis of COL6 and marker genes, annotation to predefined cell types and dot blot generation were performed via the CELL×GENE gene expression tool. An analysis of the expressions of *COL6A1*, *COL6A2*, *COL6A3, SERPINE1* and *SERPINE2* in distinct cell populations in different cancers was conducted using the Curated Cancer Cell Atlas as a resource^68^. The Curated Cancer Cell Atlas is based on a large database of scRNA sequencing datasets.

### Spatial Transcriptomics

Spatial transcriptomics of ccRCC tissue (Visium platform, spot size 55 µm in diameter, 10x Genomics) was analyzed using a previously published dataset of a cohort of 5 patients^69^, downloaded as h5ad object from Mendeley Data (https://doi.org/10.17632/g67bkbnhhg.1). The processed data only from the “tumor core” was used and reassessed with Scanpy (v.1.10.2). Gene scores were computed using the “score_genes” tool from Scanpy. The COL6 Score was computed from the *COL6A1*, *COL6A2* and *COL6A3* genes. The proliferative score was computed based on previously described cell cycle genes^70^. The SerpinE score was computed based on *SERPINE1* and *SERPINE2* gene expression. A co-occurrence score was computed by converting each gene signature score to percentile ranks across spots and multiplying these normalized ranks per spot to quantify their joint enrichment. Furthermore, a Spearman‘s correlation was performed based on the gene scores of each Visium spot.

### Transcriptome analysis of cell- and tissue culture

RNA sequencing was performed as recently described^71^. In brief equalized numbers of respective cell lines were seeded in 10 cm dishes and cultured in 10 ml culture medium (or respectively supplemented) at 37°C, 5% CO_2_ in a humidified incubator for 24 h. Cells were harvested by scraping and snap freezing before proceeding with RNA isolation. For TK173 and 786-O, comprising shCTRL and shCOL6 RNA, 3 independent replicates per condition were isolated. For A498 reseeded on shCTRL- vs shCOL6-CDMs, 3x10^6^ cells were seeded on previously prepared respective CDMs as described above, following incubation for 24 hours. RNA of 2 independent replicates per condition was isolated. For TK173, DMSO vs cabozantinib cells were treated during the 24 h incubation with 0.1% DMSO or 5 µM cabozantinib, respectively. RNA of 3 independent replicates per condition was isolated. Isolation of total RNA was performed using the Monarch Total RNA Miniprep Kit (T2010S, New England Biolabs Inc.) in accordance with the supplier’s protocol (including optional DNase-I digest). RNA quality was assessed using RNA quality numbers (RQN, Agilent Fragment Analyzer), ranging from 8.9 to 10. Poly(A) mRNA selection, library preparation and Illumina 150 pb paired-end sequencing were performed by GENEWIZ (GENEWIZ Germany GmbH, Leipzig, Germany). The Galaxy Europe platform^72^ was used for analysis of transcriptome data (applying fastp v0.23.2 or v0.24.0, HISAT2 v2.2.1, featureCounts v2.0.1 or v2.0.8 and DESeq2 v2.11.40.8 tools).

Slice culture of ccRCC samples of 5 patients was performed as described above. Samples were FFPE-embedded, 10 µm sections were generated, deparaffinized and total RNA was isolated using the Qiagen RNAeasy FFPE Kit (73504, QIAGEN GmbH, Germany). rRNA depletion, library preparation and Illumina 150 pb paired-end sequencing were performed by GENEWIZ (GENEWIZ Germany GmbH, Leipzig, Germany). The Galaxy Europe platform^72^ was used for analysis of transcriptome data (applying fastp v0.23.2, HISAT2 v 2.2.1, featureCounts v2.0.1 and DESeq2 2.11.40.8 tools). For differential gene expression analysis by DEseq2 a two-factor analysis was applied, accounting for matching treated samples from the same patient.

Gene Ontology (GO) enrichment analysis was carried out using clusterProfiler with Gene Ontology databases^73^. Extracellular matrix proteins were classified using MatrisomeAnalyzeR (v1.0.1)^74^. A list of immunomodulatory genes was derived from a previously published pan-cancer analysis of The Cancer Genome Atlas Research Network^75^.

Results of transcriptome analysis are available in Supplementary Spreadsheet S2 and are accessible via GEO series, accession numbers GSE287609, GSE287610 and GSE287677.

### Proteomic analysis of CDMs

CDM samples for proteomic analysis were generated as described above. Samples were mixed with 2x S-Trap lysis buffer and urea to final concentrations of 5 % (m/v) SDS, 50 mM triethylammonium bicarbonate (TEAB) and 6M urea at pH 8.5. Sample were sonicated in a Bioruptor Plus (Diagenode, Belgium; high energy, 20 cycles, 40 s on / 20 s off), heated (10 min, 70 °C, 500 rpm), and centrifuged (10 min, 20,800 rcf). Protein concentrations were estimated using the BCA assay (Thermo Fisher Scientific), followed by protein reduction (5 mM TCEP) and alkylation (20 mM CAA) for 30 min at 37 °C in the dark. Samples were prepared, loaded onto S-Trap Micro Spin columns (Protifi, USA) and washed according to the manufacturer’s protocol. Further, samples were treated with 1000 u PNGase F (Serva) for 30 min at 37 °C and washed twice with S-Trap binding buffer. Proteins were digested using a protease mix comprising Lys-C (Serva) and trypsin (Serva) at a 1:100 and 1:25 protease:protein ratio, respectively, and incubated overnight at 37 °C. Peptides were eluted according to the manufacturer’s protocol and dried. After reconstitution in H2O, peptide concentrations were estimated using the peptide BCA assay (Thermo Fisher Scientific).

For LC-MS/MS analysis, 500 ng of peptides were spiked with 100 fmol iRTs and loaded on Evotips (Evosep, Denmark) according to the manufacturer’s protocol. Peptides were separated using the 30SPD method (44 min gradient, 500 nL/min) on an Evosep One chromatography system (Evosep) with a 15 cm reversed-phase EV1137 performance column (Evosep). Buffer A consisted of 0.1 % formic acid in water and buffer B of 0.1 % formic acid in ACN. Mass spectrometry analysis was carried out using a timsTOF flex mass spectrometer (Bruker Daltonics, Germany) equipped with a CaptiveSpray ion source. The mass spectrometer operated in positive dia-PASEF mode with an MS1 m/z range of 100–1700, MS2 m/z range of 300–1200 and an ion mobility (1/K0) range of 0.7–1.3 V*s/cm2. Collision energies were set to 20–59 eV and ion accumulation time to 100 ms with 100 % duty cycle. The capillary voltage was set to 1600 V and drying gas temperature to 3 L/min at 180 °C. DIA acquisition employed 32 optimized isolation windows across the m/z range, resulting in a total cycle time of 1.80 s.

Raw LC-MS/MS data were processed using DIA-NN (v1.9.2)^76, 77^. A predicted spectral library was generated from the human reference proteome (downloaded from https://www.ebi.ac.uk/reference_proteomes/ on 22/07/2023), appended with common contaminants and 11 iRT peptides. The match-between-runs (MBR) algorithm was enabled to refine the predicted library. The search included peptide precursors with lengths of 7–30 amino acids, charge states of 1–4, and an m/z range of 100–1700, allowing one missed cleavage. Fragment ion m/z values were restricted to 290–1210, and identifications were filtered at a 1 % false discovery rate (FDR). Results of CDM proteome analysis are available in Supplementary Spreadsheet S3.

### Proteomic analysis of ccRCC cohort samples

Tissue specimens were collected between 2009 and 2022 during surgical tumor resection at the University Medical Center Freiburg. After resection, tissues were transferred into formalin solution and sectioned, processed, and analyzed following standard protocols for histopathological diagnostics, including embedding in paraffin. For proteomic analysis, 10 μm thick tissue slices were automatically deparaffinized and dissected into tumor and NAT compartments guided by corresponding H&E staining by an experienced pathologist. Subsequent sample preparation, LC-MS/MS analysis (Q Exactive Plus mass spectrometer, Thermo Fisher Scientific) and data-independent acquisition (DIA) proteomics analysis were performed as described previously^6^. Results of ccRCC proteome analysis are available in Supplementary Spreadsheet S4.

### Quantification and statistical analysis

Statistical analysis and graphical depiction of experimental data were performed using R (version 4.5.1) and RStudio (version 2024.12.1) as well as GraphPad Prism 8 and 10. Dots in plots indicate the respective individual units (independent experiments, replicates, samples, patients) used for statistical testing, while bars or single dots indicate mean values of replicates or samples, described accordingly in detail within the respective figure legend. Error bars indicate the standard error of the mean (S.E.M.). Bars in violin plots indicate median and quartiles, while dots indicate individual cells or measurements.

Statistical testing was performed in accordance with experimental design (two-sided, multiple comparison, correlation analysis, survival analysis) and data distribution. Data were tested for parametric distribution (Shapiro-Wilk test) and unequal variance (Levene’s test, F test) and tests were applied accordingly. Analysis of proteomics and transcriptomics datasets was performed as described in the respective methods sections. Statistical significance was defined as p < 0.05 and significance levels are indicated as ∗ – p < 0.05, ∗∗ – p < 0.01, ∗∗∗ – p < 0.001, ∗∗∗∗ – p < 0.0001 and non-significant (n. s.) in the respective figure panels. The number of independent experiments and analyzed units are stated in the figure legends and/or respective methods section.

## Acknowledgements

We thank Katja Gräwe, Alena Sammarco, Marlene Schmid, Mandana Noodeh, Severine Kayser and Heike Herzog, for expert technical assistance. In addition, we would like to express our gratitude to all members of our laboratories for their helpful discussions and support. The authors acknowledge the support of the Freiburg Galaxy Team: Björn Grüning, Bioinformatics, University of Freiburg (Germany), funded by the Collaborative Research Centre 992 Medical Epigenetics (DFG grant SFB 992/1 2012) and the German Federal Ministry of Education and Research BMBF grant 031 A538A de.NBI-RBC. We thank the Lighthouse Core Facility staff of the Medical Center - University of Freiburg for their help with their resources and their excellent support. The Lighthouse Core Facility is funded by the Medical Faculty, University of Freiburg (Project Numbers 2021/A2-Fol; 2021/B3-Fol). The atomic force microscope, NanoWizard III (JPK, Bruker nano) was partly funded by the Ministry of Science, Research and the Arts of Baden-Württemberg (Az: 33-7532.20) and the University of Freiburg (“Strategiefonds”), granted to Winfried Römer.

## Author contributions

Conceptualization, MR and CS; Methodology, MW GA, CLE, WR, TV, MH, OS, MR and CHS; Formal analysis, MW, GA, TF, CGP, MR, OS and CHS; Investigation, MW, GA, TF, ND, MZ, CGP, ALK, CH, AN, MH, TV and MR; Resources, CLE, TV and MG; Writing – original draft preparation, MW, GA, MR and CS; Writing – review and editing, MW, GA, MR and CS, with input from all authors; Visualization, MW, GA, MR and CS; Supervision, MR and CS; Project administration, MW, OS, MR and CS; Funding acquisition, WR, OS and CS. Parts of this work were part of the doctoral theses of MW and CGP. All authors have read and agreed to the published version of the manuscript.

## Funding

This work was supported by the following grants: Deutsche Forschungsgemeinschaft (DFG, German Research Foundation) – grant 431984000 (SFB 1453), (C.S. and O.S.); grant 441891347 (SFB 1479); grant 241702976, DFG SCHE2092/3-1 (C.S.); grant 438496892, DFG SCHE2092/4-1 (C.S.); grant 256073931 (SFB 1160), (C.S.); grant 501370692, DFG SCHE2092/5-1 (C.S.); grant 562400794, DFG SCHE2092/6-1 (C.S.); grant 559178177 (INST 39/1466-1); grant 241702976, DFG SCHE 2092/1-3; and Wilhelm Sander-Foundation – grant 2023.010.1 (C.S.). C.L.E. and W.R. acknowledge funding by the Deutsche Forschungsgemeinschaft under Germany’s Excellence Strategy (CIBSS - EXC-2189 - Project ID 390939984).

## Competing interests

The authors declare no competing interests.

## Data and materials availability

The data presented in this study are available in the article or supplementary materials. Previously published datasets are available from the original publication or databases as described in the respective methods sections. Resources generated in this study are available from the corresponding author upon reasonable request.

## Supplementary information

Supplementary Figure S1. Extended data related to main Figure 1.

Supplementary Figure S2. Extended data related to main Figure 2.

Supplementary Figure S3. Extended data related to main Figure 3.

Supplementary Figure S4. Extended data related to main Figure 4.

Supplementary Figure S5. Extended data related to main Figure 5.

Supplementary Figure S6. Extended data related to main Figure 6.

Supplementary Figure S7. Extended data related to main Figure 7.

Supplementary Spreadsheet-S1: Antibodies and staining information.

Supplementary Spreadsheet-S2: RNA sequencing analysis.

Supplementary Spreadsheet-S3: Proteomics analysis of CDMs.

Supplementary Spreadsheet-S4: Proteomics analysis of ccRCC samples.

## Notes

### Competing Interest Statement

The authors have declared no competing interest.

