## Supplementary Figures for "Fibroblast-derived Collagen VI shapes the structure and function of the tumor-immune microenvironment in clear cell renal cell carcinoma"

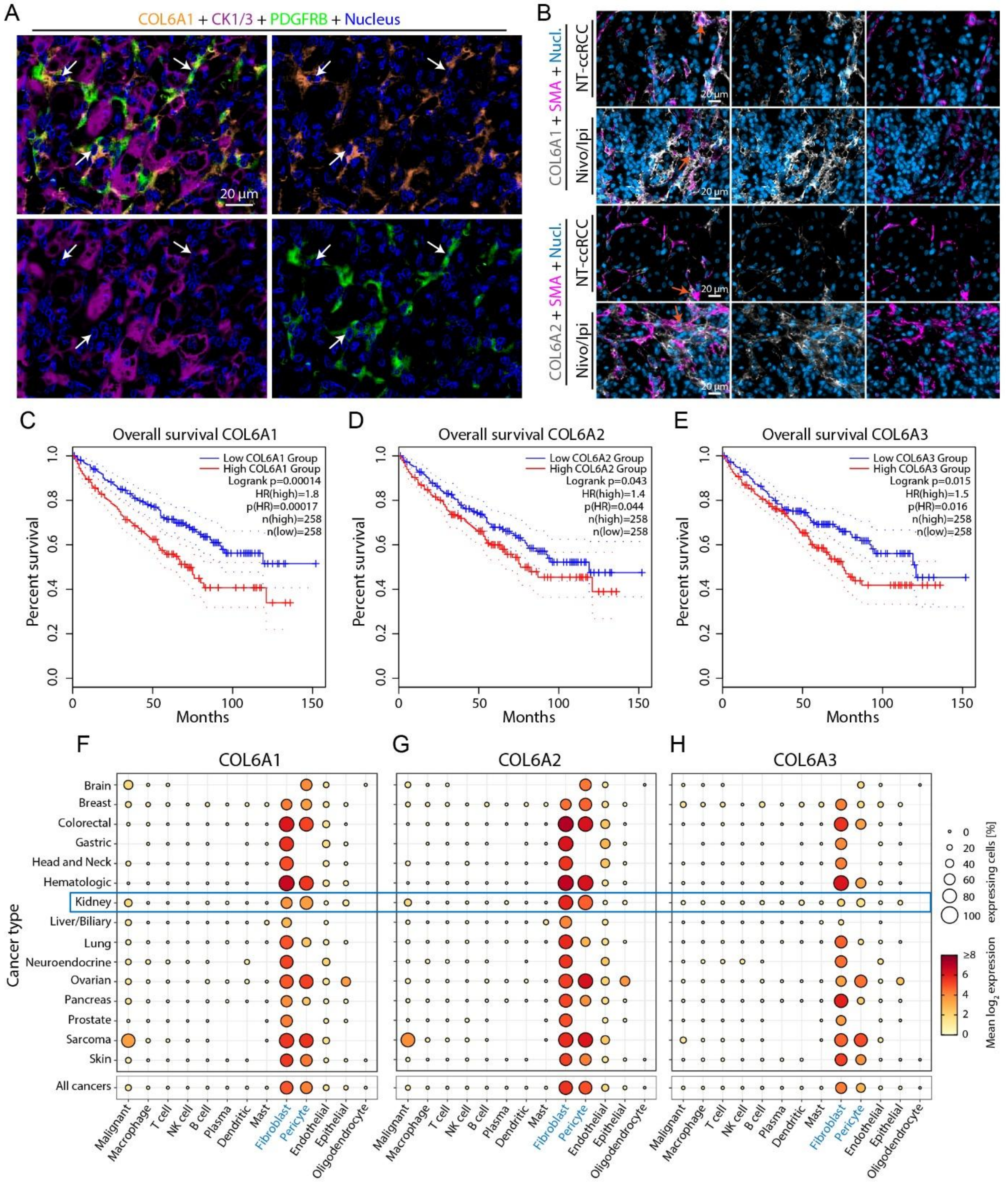


**Supplementary Figure S1. Extended data related to main Figure 1**

**(A)** Multiplex IF staining indicates partial expression of COL6A1 (orange) by PDGFRB (green) positive cells (white arrows) in ccRCC. Co-staining for CK (purple) and DNA (blue). **(B)** Multiplex IF staining showing similar expression patterns of COL6A2 (gray) in ccRCC as for COL6A1 staining, further demonstrating partial expression of COL6A2 by SMA (violet) positive cells (orange arrows), co-staining for DNA (blue). Analysis of a ccRCC case treated with immune checkpoint inhibitors Nivolumab and Ipilimumab indicates increased expression of COL6A2. **(C-E)** Survival analysis for high or low expression of *COL6A1*, *COL6A2* and *COL6A3* in the TCGA ccRCC cohort. Graphs show Kaplan-Meier plots and Log-rank tests. **(F-G)** scRNA sequencing analysis of *COL6A1*, *COL6A2* and *COL6A3* expression in the Curated Cancer Cell Atlas.

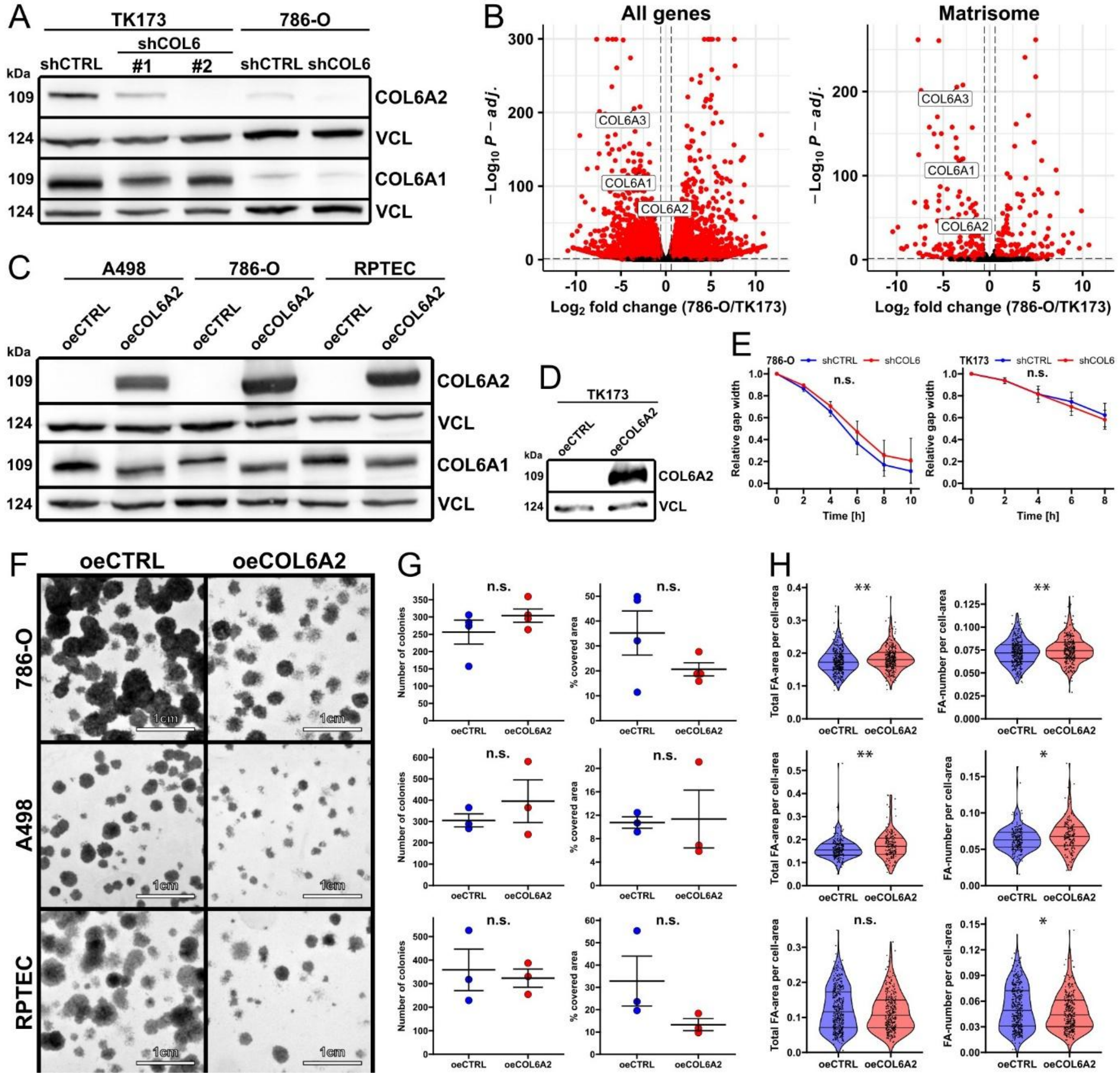


**Supplementary Figure S2. Extended data related to main Figure 2**

**(A)** Validation of the *COL6A2* knockdown in TK173 and 786-O cell lines. Western blot for COL6A1, COL6A1. For TK173, two independent shRNA clones were evaluated. **(B)** Volcano plot depicting differentially expressed genes between 786-O shCTRL and TK173 shCTRL cell lines (red dots indicate significantly regulated genes) **(C)** Western blot validation of overexpression of *COL6A2* in A498, 786-O and RPTEC cell lines for COL6A1 and COL6A2. **(D)** Validation of *COL6A2* overexpression in the TK173 cell line. Western blot for COL6A2. **(E)** Line plots showing gap closure assays with relative gap width from the starting point over time of shCTRL (blue) and shCOL6 (red) for 786-O (left) and TK173 (right) cell lines (N=3 independent experiments). **(F-G)** Representative images and quantification analysis of colony formation assay of oeCTRL and oeCOL6A2 in 786-O (top), A498 (middle) and RPTEC (bottom) cell lines (N=3 independent experiments, for 786-O N=4 independent experiments, unpaired t test). **(H)** FA quantification depicted with violin plots of oeCTRL and oeCOL6A2 in 786-O (top), A498 (middle) and RPTEC (bottom) cell lines (dots indicate cells from 3 independent experiments, Mann-Whitney U test). Bars indicate mean and S.E.M in dot plots and line plots, or median and quartiles in violin plots; $*$ – $p < 0.05$, $**$ – $p < 0.01$ and n.s – not significant.

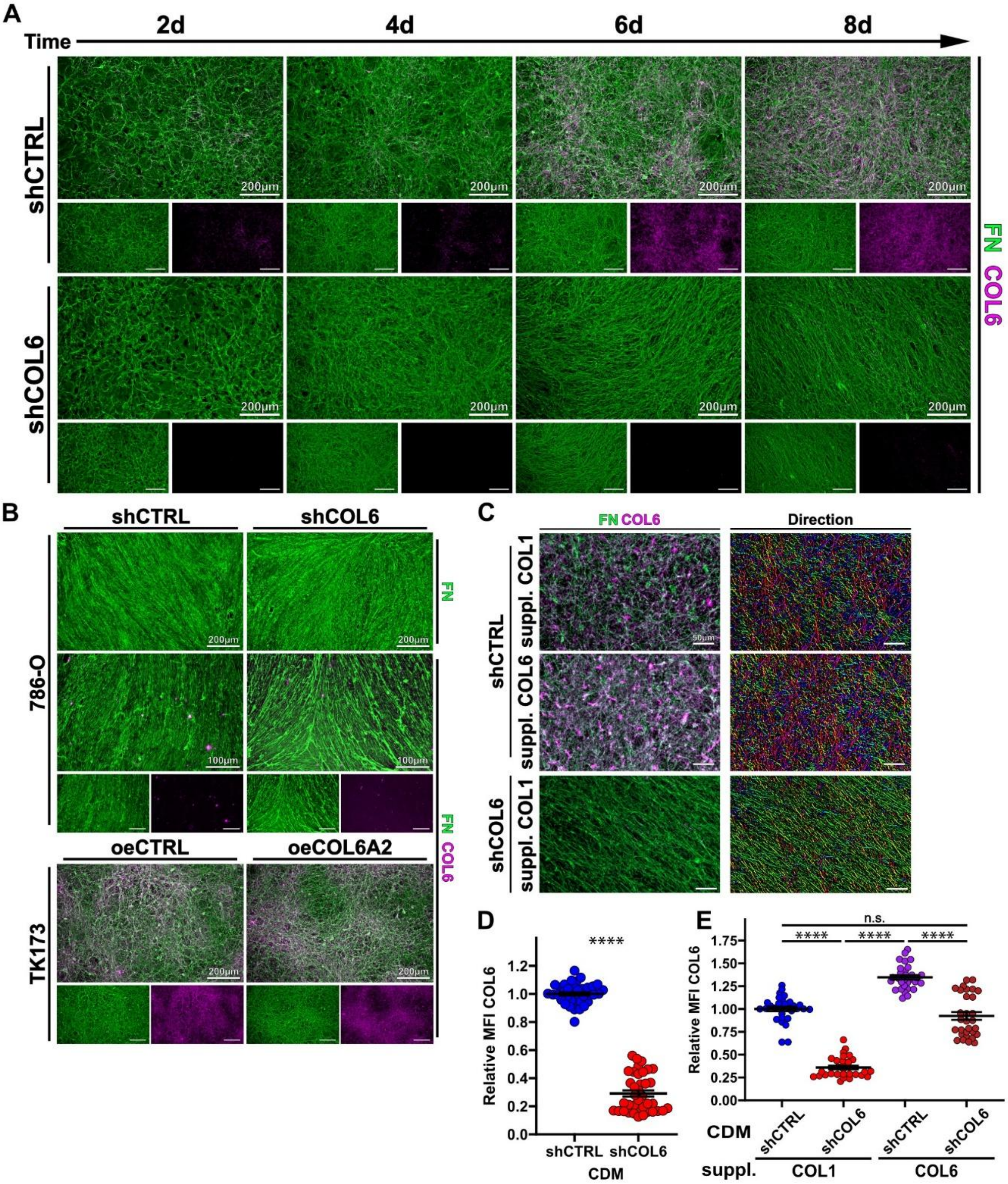


**Supplementary Figure S3. Extended data related to main Figure 3**

**(A-C)** IF images stained for FN (green) and COL6 (violet) of different CDMs derived from different cell backgrounds. **(A)** CDM and ECM composition from TK173 shCTRL and shCOL6 over time (2, 4, 6 and 8 days). **(B)** CDM from 786-O shCTRL and shCOL6 (top). CDM from TK173 oeCTRL and oeCOL6A2. **(C)** CDMs from TK173 shCTRL supplemented with COL1 and COL6, as well as shCOL6 supplemented with COL1. Color-coded FN fiber directionality in the right panel. **(D)** Quantification of the normalized mean fluorescence intensity (MFI) of COL6 between shCTRL and shCOL6 (N=37 shCTRL and 38 shCOL6 images pooled from 3 independent experiments, Mann-Whitney U test). **(E)** Dot plot depicting normalized MFI of shCTRL and shCOL6 CDMs supplemented with COL6 and COL1 (30 images pooled from 3 independent experiments, Kruskal-Wallis test, followed by Dunn's post hoc test with Holm-Bonferroni correction). Bars indicate mean and S.E.M in dot plots; **** – $p < 0.0001$ and n.s – not significant.

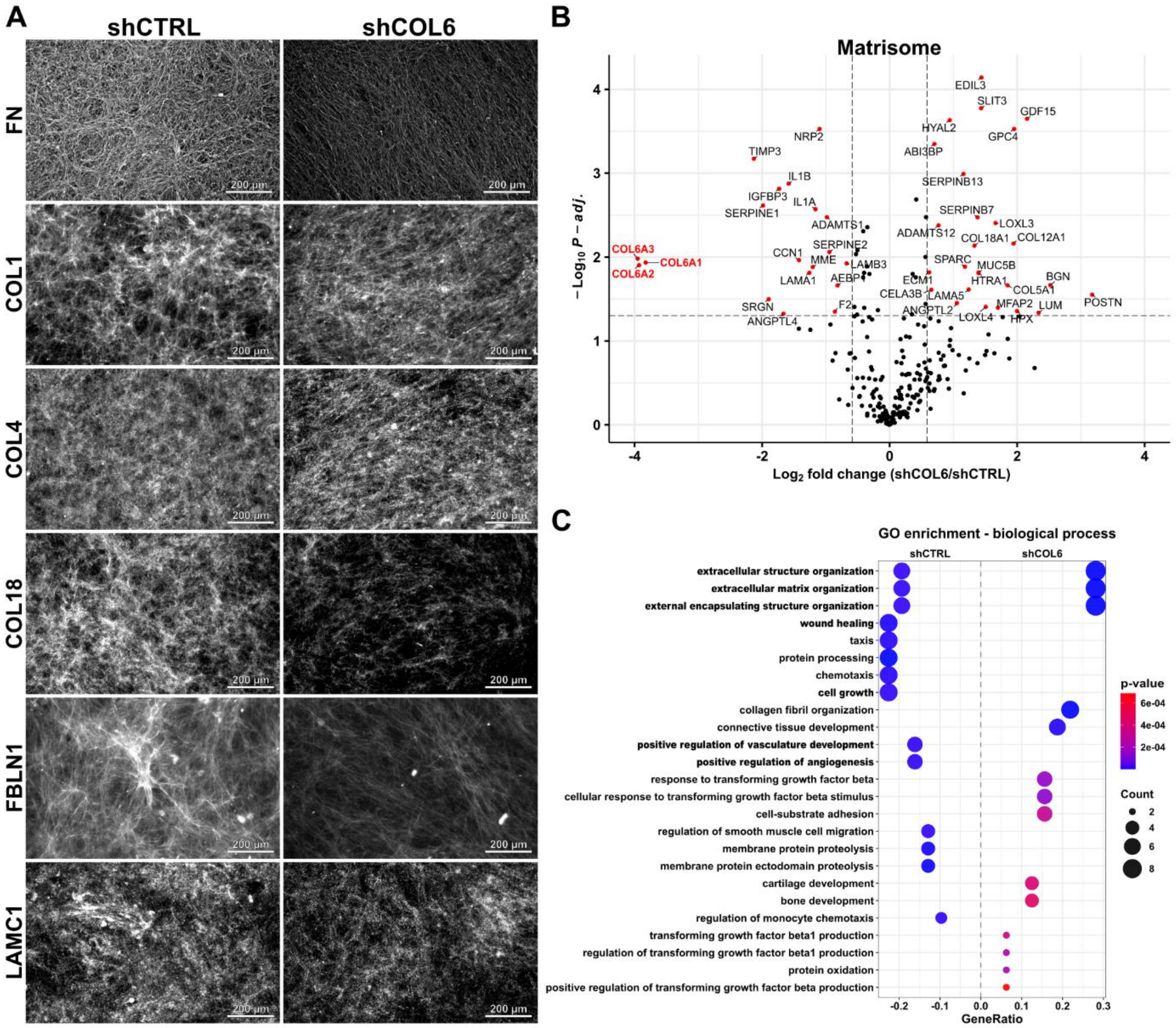


**Supplementary Figure S4. Extended data related to main Figure 4**

**(A)** IF staining for indicated ECM proteins (white) in CDMs from shCTRL and shCOL6 cells. **(B)** Volcano plot of CDM MS-analysis showing differentially expressed matrisome proteins in shCOL6 compared to shCTRL. **(C)** GO enrichment analysis of the biological processes of significantly regulated proteins depicted in a bubble plot.

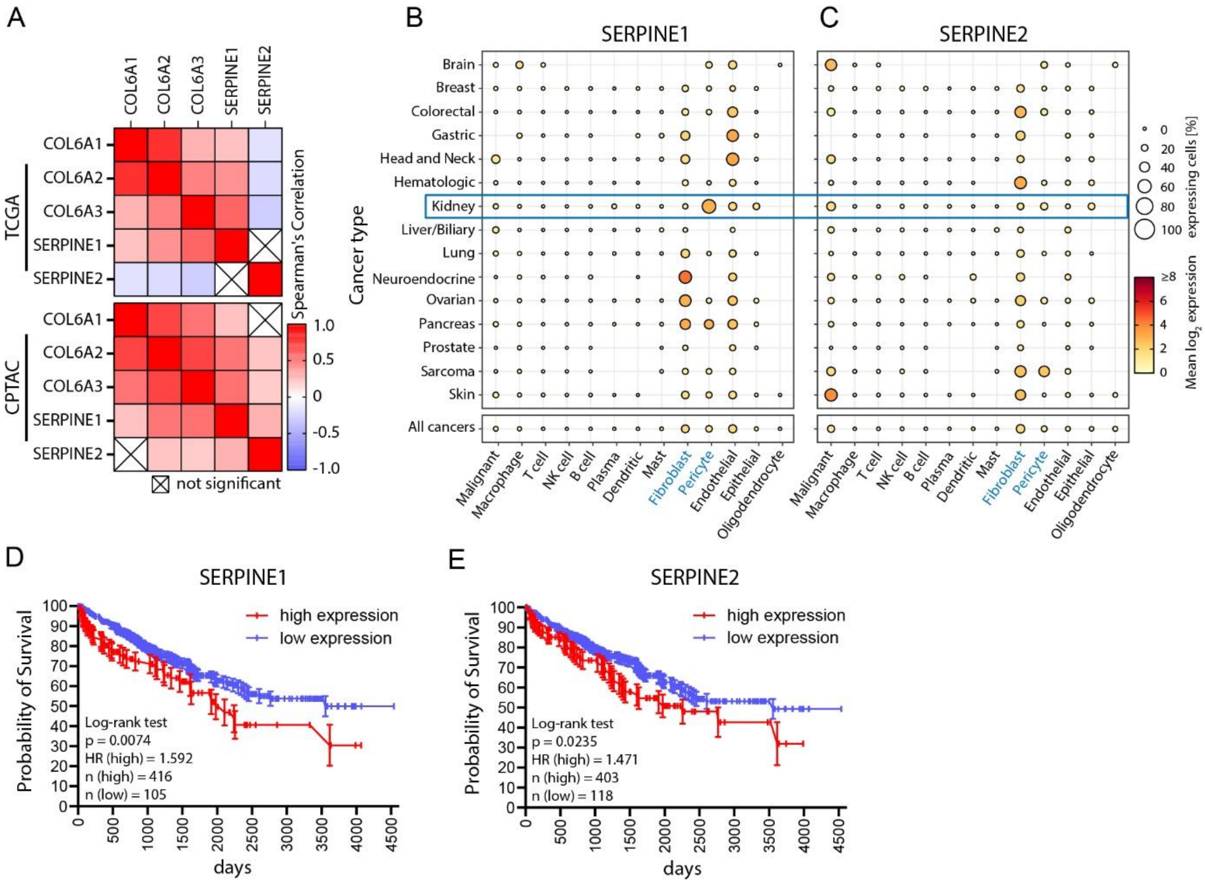


**Supplementary Figure S5. Extended data related to main Figure 5**

**(A)** Heatmap depicting Spearman's correlation analysis of *COL6A1*, *COL6A2, COL6A3*, *SERPINE1* and *SERPINE2* gene expression in the ccRCC TCGA and CPTAC cohorts. **(B&C)** scRNA sequencing analysis of *SERPINE1* and *SERPINE2* expression in the Curated Cancer Cell Atlas. **(D&E)** Survival analysis for high or low expression of *SERPINE1* and *SERPINE2* in the TCGA ccRCC cohort. Graphs show Kaplan-Meier plots and Log-rank (Mantel-Cox) tests.

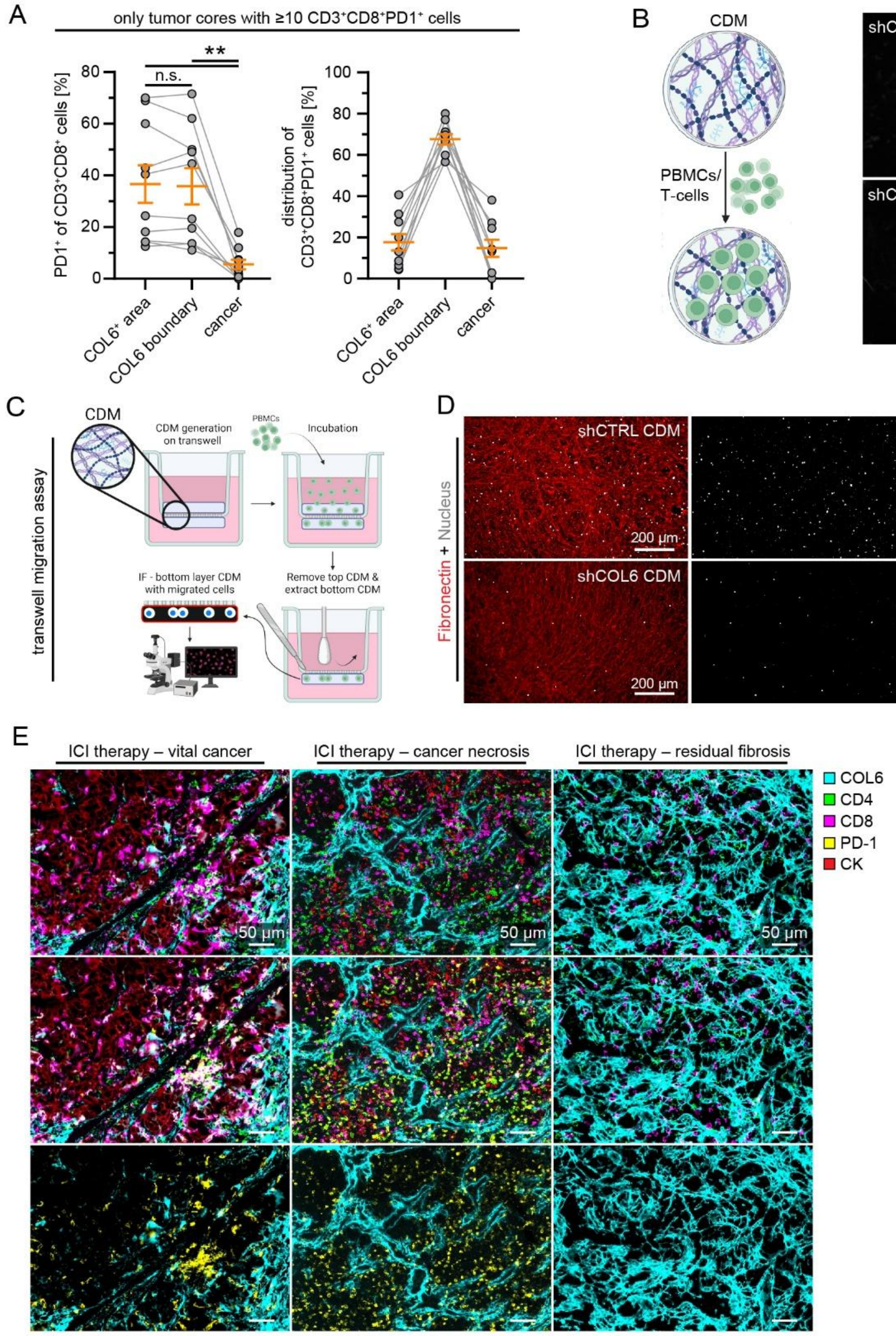


**Supplementary Figure S6. Extended data related to main Figure 6**

**(A&B)** SeqIF analysis of the percentage of CD3+CD8+PD1+ cells of total CD3+CD8+ cells in indicated compartments and relative distribution of CD3+CD8+PD1+ cells to indicated compartments, excluding samples with very low numbers of CD3+CD8+PD1+ cells (dots indicate N=10 cores, bars indicate mean and S.E.M, ∗∗ – $p < 0.01$ and non-significant (n. s.), RM one-way ANOVA with Geisser-Greenhouse correction and Tukey's multiple comparison test). **(B)** Schematic description of T cell and PBMC reseeding on previously prepared CDM (Created in BioRender. Schell, C. (2026) https://BioRender.com/x5sqb3y). **(C)** Schematic description of CDM-transwell assay (Created in BioRender. Schell, C. (2026) https://BioRender.com/x5sqb3y). **(D)** IF staining of DNA (white) and FN (red) of PBMCs migrated into the transwell's bottom layer CDM. **(E)** SeqIF analysis of an immune checkpoint inhibitor (ICI) treated ccRCC patient sample. Images show overviews and single channels of images shown in main Figure 6. Representative images of regions with vital cancer, immune infiltration and cancer necrosis, and residual fibrosis are shown (markers as indicated).

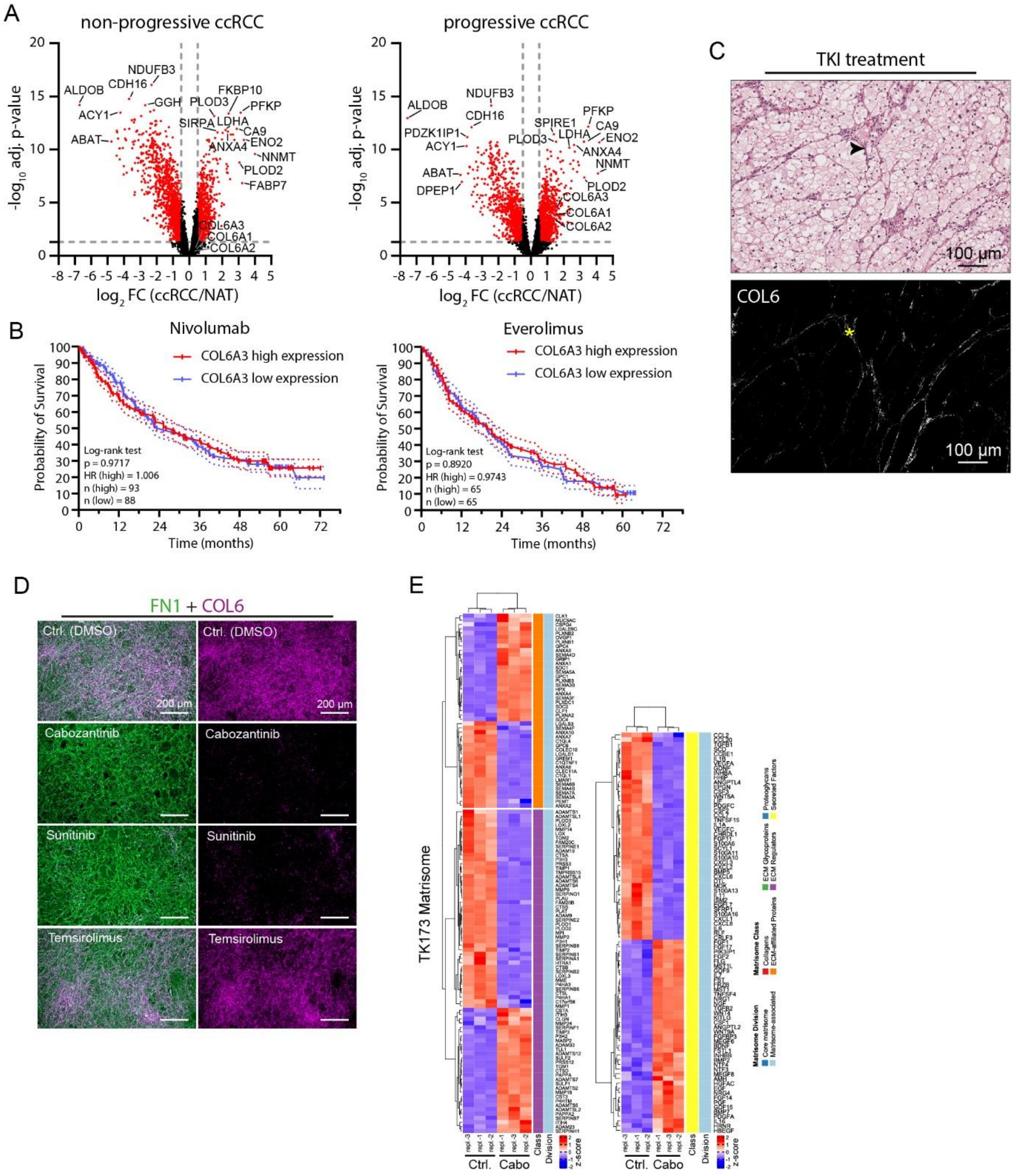


**Supplementary Figure S7. Extended data related to main Figure 7**

**(A)** Vulcano plot of ccRCC proteome analysis separated for patients with non-progressive and progressive disease. Red dots indicate proteins with adjusted p-value <0.05 and $log_2$ FC (fold change) > |0.5|; NAT – normal adjacent tissue). **(B)** Survival analysis of high or low expression of COL6A3 in the CheckMate 025 study cohort of Nivolumab (anti-PD-1) or everolimus (mTOR inhibitor) treated patients. Graphs show Kaplan-Meier plots and Log-rank (Mantel-Cox) tests. **(C)** Representative HE and COL6 IF on a serial section staining of a ccRCC patient that received neoadjuvant TKI treatment with cabozantinib (arrowhead indicate stromal septa and yellow asterisk COL6 positive septa in sequential sections). **(D)** IF images stained for FN (green) and COL6 (violet) of synthesized CDMs of TK173 cells treated with the indicated drugs. Images show overviews and single channels of images shown in main Figure 7. **(H)** Heatmap of significantly regulated matrisome genes in cabozantinib-treated TK173 cells (genes with adjusted $p<0.0001$ and fold change > |2| are depicted).
